# A genetically encoded fluorescent biosensor for extracellular L-lactate

**DOI:** 10.1101/2021.03.05.434048

**Authors:** Yusuke Nasu, Ciaran Murphy-Royal, Yurong Wen, Jordan Haidey, M. Rosana S. Molina, Abhi Aggarwal, Shuce Zhang, Yuki Kamijo, Marie-Eve Paquet, Kaspar Podgorski, Mikhail Drobizhev, Jaideep S. Bains, M. Joanne Lemieux, Grant R. Gordon, Robert E. Campbell

## Abstract

To enable investigations of the emerging roles of cell-to-cell shuttling of L-lactate, we have developed an intensiometric green fluorescent genetically encoded biosensor for extracellular L-lactate. We demonstrate that this biosensor, designated eLACCO1.1, enables minimally invasive cellular resolution imaging of extracellular L-lactate in cultured mammalian cells and brain tissue.

## Introduction

Organisms on Earth produce biologically-available energy by catabolizing glucose to pyruvate which, in the presence of oxygen, is normally catabolized to carbon dioxide or, in the absence of oxygen, is normally converted into L-lactate. Traditionally, L-lactate (the anionic form that predominates at physiological pH) and L-lactic acid (the protonated form that predominates at acidic pH) have been considered “waste” by-products of glucose metabolism^1^. However, growing evidence suggests that L-lactate is better considered a biological “fuel currency”, that can be shuttled from cell-to-cell and plays a central role in the energy supply for organisms^1^. For example, the astrocyte-to-neuron lactate shuttle (ANLS) hypothesis proposes that astrocytes metabolize glucose to produce L-lactate which is then released to the extracellular environment and taken up by neurons. In neurons, L-lactate is converted to pyruvate which is fed into the tricarboxylic acid cycle (TCA) for production of adenosine triphosphate (ATP) which provides energy necessary to sustain heightened neural activity^2^. The ANLS hypothesis remains somewhat controversial, with recent reports of evidence both for^2^ and against^3^ it.

Investigations of cell-to-cell L-lactate shuttles, and testing of the ANLS hypothesis in particular, would be greatly facilitated by the availability of a genetically encoded fluorescent biosensor that would enable high resolution spatially and temporally resolved imaging of changes in the concentration of extracellular L-lactate. Genetically encoded fluorescent biosensors are powerful tools for cell-based and *in vivo* imaging of molecules, ions, and protein activities, and many examples have been reported to date^4, 5^. However, despite the remarkable progress in the development of such biosensors, including ones for intracellular L-lactate^6, 7^, no genetically encoded biosensors for extracellular L-lactate have yet been reported.

## Results

### Development and characterization of a genetically encoded L-lactate biosensor, eLACCO1

To construct an initial prototype L-lactate biosensor, we inserted circularly permuted green fluorescent protein (cpGFP) into *Thermus thermophilis* TTHA0766 L-lactate binding protein at 70 different positions. Positions were chosen by manual inspection of the protein crystal structure to identify regions that were solvent exposed and likely to undergo L-lactate dependent conformational changes (**Supplementary Fig. 1**)^8^. The insertion variant with the largest change in fluorescence intensity (Δ*F*/*F* = (*F*_max_ - *F*_min_)/*F*_min_) upon L-lactate treatment (designated eLACCO0.1) had cpGFP inserted at position 191, and exhibited an inverse response (fluorescence decrease upon binding) with Δ*F*/*F* = 0.3. Efforts to create prototype biosensors by inserting cpGFP into the analogous regions of TTHA0766 homologues produced no variants with larger L-lactate-dependent changes in fluorescence intensity (**Supplementary Fig. 2**). Accordingly, we focused our efforts on further development of eLACCO0.1.

To develop variants of eLACCO0.1 with larger Δ*F*/*F*, we performed directed evolution with screening for L-lactate-dependent change in fluorescence intensity (**Supplementary Fig. 3a**). The first round of evolution used a library in which residues of the C-terminal linker were randomized. Screening of this library led to the discovery of eLACCO0.2 with a direct response (fluorescence increase upon binding) to L-lactate (Δ*F*/*F* = 0.3; **Supplementary Fig. 3b**). Additional rounds of evolution by random mutagenesis of the entire biosensor gene ultimately produced the eLACCO1 variant with Δ*F*/*F* of 6 and excitation and emission peaks at 494 and 512 nm, respectively (**Fig. 1a** and **Supplementary Figs. 3c,d and 4**). The crystal structure of TTHA0766 reveals an integral and essential calcium ion (Ca^2+^) in the L-lactate binding pocket^8^, indicating that L-lactate binding should be Ca^2+^ dependent. To investigate the effect of Ca^2+^ on the biosensor functionality, we measured the fluorescence intensity of eLACCO1 in the presence of L-lactate and Ca^2+^. These experiments revealed that Ca^2+^ is indeed essential for the L-lactate-dependent response of eLACCO1, and the biosensor only functions as an L-lactate biosensor at concentrations of Ca^2+^ greater than 0.6 μM (**Supplementary Fig. 5a,b**). eLACCO1 showed a pH dependence similar to that of the GCaMP6f Ca^2+^ biosensor, with p*K*_a_ values of 6.0 and 8.7 in the presence and absence of L-lactate, respectively (**Supplementary Fig. 5c**)^9^. Determination of L-lactate-binding affinity with L-lactate titration revealed that an apparent dissociation constant (*K*_d_) of eLACCO1 is 4.1 and 120 μM for L-lactate and D-lactate, respectively (**Supplementary Fig. 5d**). Relative to L-lactate, eLACCO1 was ∼320, ∼460 and ∼1320 times less sensitive to the structurally similar molecules β-hydroxybutyrate, pyruvate and oxaloacetate, respectively (**Supplementary Fig. 5d**). Negligible responses were observed for all other metabolites tested.

**Fig. 1.**
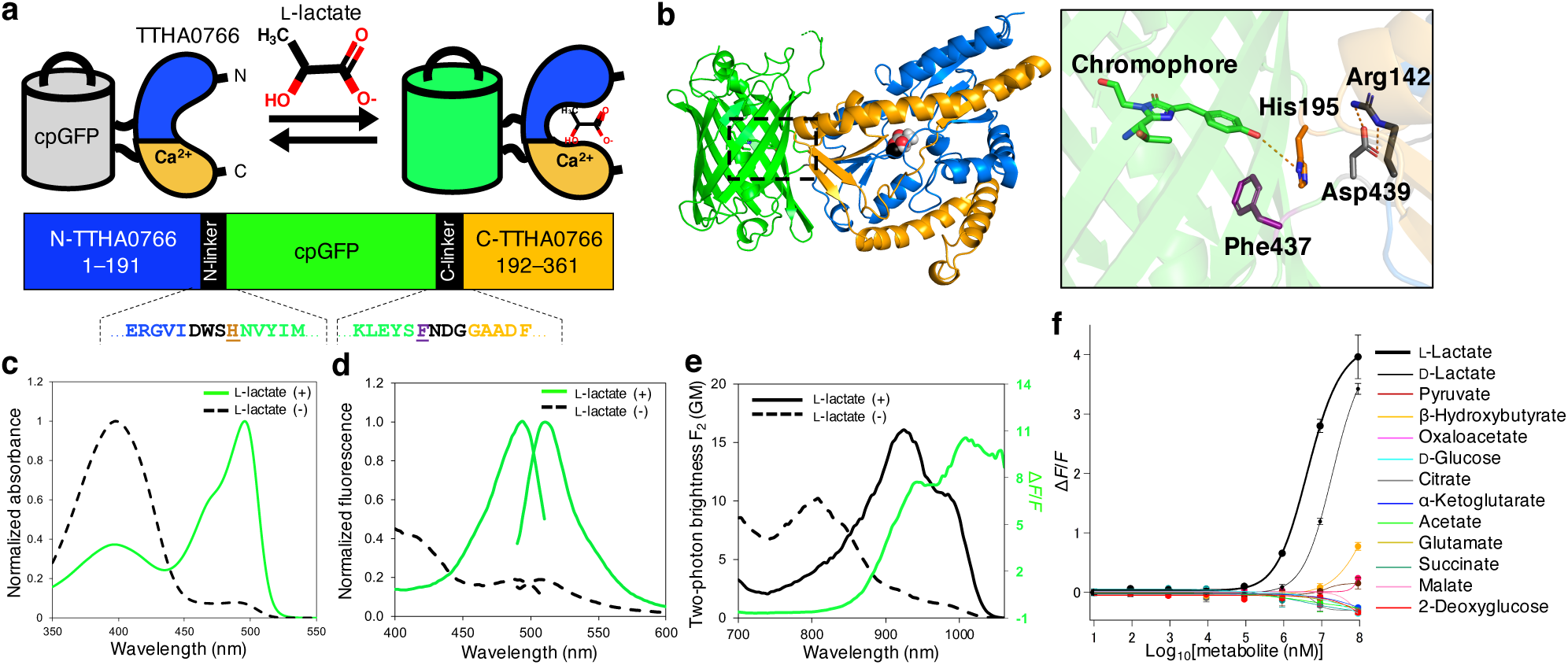
Structure and *in vitro* characterization of eLACCO1.1. (**a**) Schematic representation of eLACCO1 and its mechanism of response to L-lactate. Linker regions are shown in black and the two “gate post” residues^5^ in cpGFP are highlighted in dark orange (His195) and purple (Phe437). (**b**) Crystal structure of eLACCO1. Right panel represents a zoom-in view around the chromophore. (**c**) Absorbance spectra of eLACCO1.1 in the presence (100 mM) and absence of L-lactate. (**d**) Excitation and emission spectra of eLACCO1.1 in the presence (100 mM) and absence of L-lactate. (**e**) Two-photon excitation spectra of eLACCO1.1 in the presence (100 mM) and absence of L-lactate. Δ*F*/*F* is represented in the green solid line. GM, Goeppert-Mayer units. (**f**) Dose-response curves of eLACCO1.1 for L-lactate and a variety of metabolites. *n* = 3 independent experiments (mean ± s.d.).

### The X-ray crystal structure of eLACCO1

In an effort to obtain molecular insight into the structure and mechanism of eLACCO1, we determined the crystal structure of eLACCO1 in the L-lactate-bound state at a resolution of 2.25 Å (**Fig. 1b** and **Supplementary Table 1**). The overall structure reveals that the cpGFP-derived and TTHA0766-derived domains are closely associated via an extensive interaction surface that contains numerous molecular contacts. The TTHA0766-derived domain of eLACCO1 retains the same dimeric structure as TTHA0766 itself (**Supplementary Fig. 6a**)^8^. The Hill coefficient of eLACCO1 is close to one, suggesting that the protomers in the dimer do not interact cooperatively (**Supplementary Fig. 5d**). Trp509 in the dimer interface forms a hydrophobic interaction with its symmetry-related self via π-π stacking. Mutagenesis of this residue abrogated L-lactate-dependent fluorescence presumably due to disruption of the dimeric structure, providing support for the conclusion that eLACCO1 functions as a dimer.

The mechanism of eLACCO1 must involve changes in the chromophore environment that are induced by conformational changes that accompany binding of L-lactate to the TTHA0766-derived domain^5^. Based on the pH dependence, these changes in the chromophore environment are stabilizing the brightly fluorescent deprotonated form of the chromophore. The structure reveals that the imidazole side chain of His195 is likely a key moiety for mediating the chromophore protonation state (**Fig. 1b**). Further insight into the mechanism comes from examining the position of beneficial mutations discovered during biosensor optimization. In the first round of the directed evolution for optimization of C-terminal linker, the introduction of the Asn439Asp mutation converted the biosensor from having an inverse response to having a direct response (**Supplementary Fig. 3b**). Accordingly, we propose that the mechanism of eLACCO1 involves a conformational switch between two states: the L-lactate-free state where the protonated (dark) chromophore is stabilized by interaction with carboxylate side chain of Asp439 (and His195 is further away); and the L-lactate-bound state where the deprotonated chromophore (bright) is stabilized by interaction with the imidazole side chain of His195 (and Asp439 is further away). Consistent with this proposed mechanism, in the final round of the evolution, we discovered the Lys142Arg mutation that substantially improved Δ*F*/*F* (**Supplementary Fig. 3b**). The eLACCO1 crystal structure reveals that Arg142 forms a salt bridge with Asp439 in the L-lactate-bound state. This salt-bridge may further limit residual interaction of Asp439 with the chromophore (possibly due to increased distance or more effective charge neutralization) and thereby contribute to increased brightness of the L-lactate-bound state and a higher Δ*F*/*F*.

The crystal structure of eLACCO1 revealed that the N-terminal linker connecting cpGFP and TTHA0766 is relatively lengthy, leading us to construct several deletion variants in an effort to increase Δ*F*/*F* by bringing the two domains into closer proximity (**Supplementary Fig. 7a**). Indeed, the deletion variants exhibited higher Δ*F*/*F* with comparable apparent *K*_d_ *in vitro* (**Supplementary Fig. 7b**). Unfortunately, later experiments revealed that these variants were not efficiently trafficked to the membrane (**Supplementary Fig. 7c**) and so they were not further pursued.

### Development and characterization of affinity-tuned eLACCO1.1

The physiological concentration of extracellular L-lactate at rest is approximately 0.2 mM in the brain^10^ and 1 mM in serum^11^, suggesting that the affinity of eLACCO1 for L-lactate (apparent *K*_d_ ∼ 4.1 μM) would be too high to reveal physiologically relevant concentration changes. To tune the affinity of eLACCO1 for the imaging of extracellular L-lactate, we aimed to decrease its affinity for L-lactate using mutagenesis. To this end, we noted that the crystal structure of eLACCO1 reveals that the phenol moiety of Tyr80 forms a hydrogen bond with bound L-lactate (**Supplementary Fig. 8a,b**). Introduction of the Tyr80Phe mutation removed this hydrogen bond and produced the low-affinity eLACCO1.1 variant with apparent *K*_d_ = 3.9 mM. A non-responsive control biosensor, designated deLACCO, was engineered by incorporating the Asp444Asn mutation into eLACCO1.1 to abolish L-lactate binding (**Supplementary Fig. 8c,d**).

With the affinity-optimized eLACCO1.1 variant in hand, we undertook a detailed characterization of its spectral properties. eLACCO1.1 has two absorbance peaks at 398 nm and 496 nm, indicative of the neutral (protonated) and the anionic (deprotonated) chromophore, respectively (**Fig. 1c**). Consistent with the absorbance spectrum, eLACCO1.1 displays excitation peaks at 398 nm and 493 nm, and excitation at either peak produces an emission peak at 510 nm (**Fig. 1d**). Excitation of the neutral form at 398 nm results in fluorescence from the excited state of the anionic (at 510 nm), presumably due to excited-state proton transfer^12^.

The molecular brightness of eLACCO1.1 in the L-lactate bound state excited at 493 nm (corresponding to anionic form) is ∼80% of EGFP (**Supplementary Table 2**)^13^. Considering that only 43% of the sensor is present in the anionic form, the fluorescence brightness per one anionic chromophore is ∼70% larger than that of EGFP^14^. This enhancement can be explained by the fact that the anionic form of eLACCO1.1 in the L-lactate bound state has the extinction coefficient ∼60% higher than EGFP and the absorption peak of eLACCO1.1 is ∼60% narrower than EGFP. L-Lactate dependent decrease in the first peak and increase in the second peak yields the excitation ratio Δ*R*/*R* of 14, where Δ*R*/*R* = (*R*_493nm_ - *R*_398nm_)/*R*_398nm_ and *R* is *F*_bound_/*F*_unbound_. Under two-photon excitation (**Fig. 1e**), eLACCO1.1 also exhibited ratiometric changes in excitation with an L-lactate dependent increase of a 924 nm peak, and concomitant decrease of an 804 nm peak, corresponding to the anionic and neutral states of the chromophore, respectively. The intrinsic two-photon brightness (*F*_2_ = σ_2_ × φ × ρ, where σ_2_ is the two-photon absorption cross section) of L-lactate-bound eLACCO1.1 at 924 nm, *F*_2_ = 16 GM, is 42% of EGFP (*F*_2_ = 38 GM)^13, 14^. This is because only 43% of eLACCO1.1 is present in the anionic form when bound to L-lactate (**Supplementary Table 2**). Though moderately dimmer than GFP in terms of two-photon brightness, it is notable that eLACCO1.1 is 2.4-fold brighter than Citrine (*F*_2_ = 6.7 GM)^15^. The L-lactate induced two-photon excited fluorescence change (Δ*F*_2_/*F*_2_ = 8 −10, at 940 - 1000 nm) is about twice as large as the one-photon excited fluorescence change (Δ*F*/*F* = 4, at 480 - 510 nm) (**Fig. 1d, e**). A similar trend has been observed for some red genetically encoded Ca^2+^ biosensors^16^.

Investigation of the molecular specificity of eLACCO1.1 revealed that the decrease in binding affinity (due to the Tyr80Phe mutation) extended to other structurally similar molecules, and pyruvate and oxaloacetate caused negligible fluorescent response even at 100 mM (**Fig. 1f**). The concentration of β-hydroxybutyrate in serum is approximately 25.5 mM even under the diabetic ketoacidosis conditions^17^, suggesting that the residual sensitivity to β-hydroxybutyrate is unlikely to be problematic for *in vivo* applications of eLACCO1.1.

### Targeting of eLACCO1.1 to the extracellular environment

To exploit eLACCO1.1 as a genetically encoded fluorescent biosensor for extracellular L-lactate in a cellular milieu, we first attempted to express the protein on the surface of live mammalian cells by fusing it to an N-terminal leader sequence and a C-terminal anchor domain (**Fig. 2a**). However, the widely used combination of immunoglobulin *κ*-chain (Ig*κ*) leader sequence and platelet-derived growth factor receptor (PDGFR) transmembrane domain for cell surface expression, resulted in only intracellular expression with localization reminiscent of the nuclear membrane (**Fig. 2b**). We screened a range of leader sequences in combination with the PDGFR anchor, but did not discover any that led to robust membrane localization of PDGFR-anchored eLACCO1.1 (**Supplementary Fig. 9**). Hence, we turned to the combination of an N-terminal leader sequence and a glycosylphosphatidylinositol (GPI) anchor, both of which are derived from CD59. This GPI-based approach did lead to the desired targeting of eLACCO1.1 to the cell surface, as indicated by colocalization with cell-surface targeted mCherry red fluorescent protein (RFP) (**Fig. 2b**). When targeted to the cell surface, eLACCO1.1, with an optimized linker between eLACCO1.1 and GPI anchor, robustly increased in fluorescence intensity (Δ*F*/*F* of ∼3) upon addition of L-lactate to the imaging medium (**Supplementary Fig. 10** and **Fig. 2c,d**). The control biosensor deLACCO had similar membrane localization and, as expected, did not respond to L-lactate (**Fig. 2c,d**). We characterized cell surface-targeted eLACCO1.1 in terms of several important parameters.

**Fig. 2.**
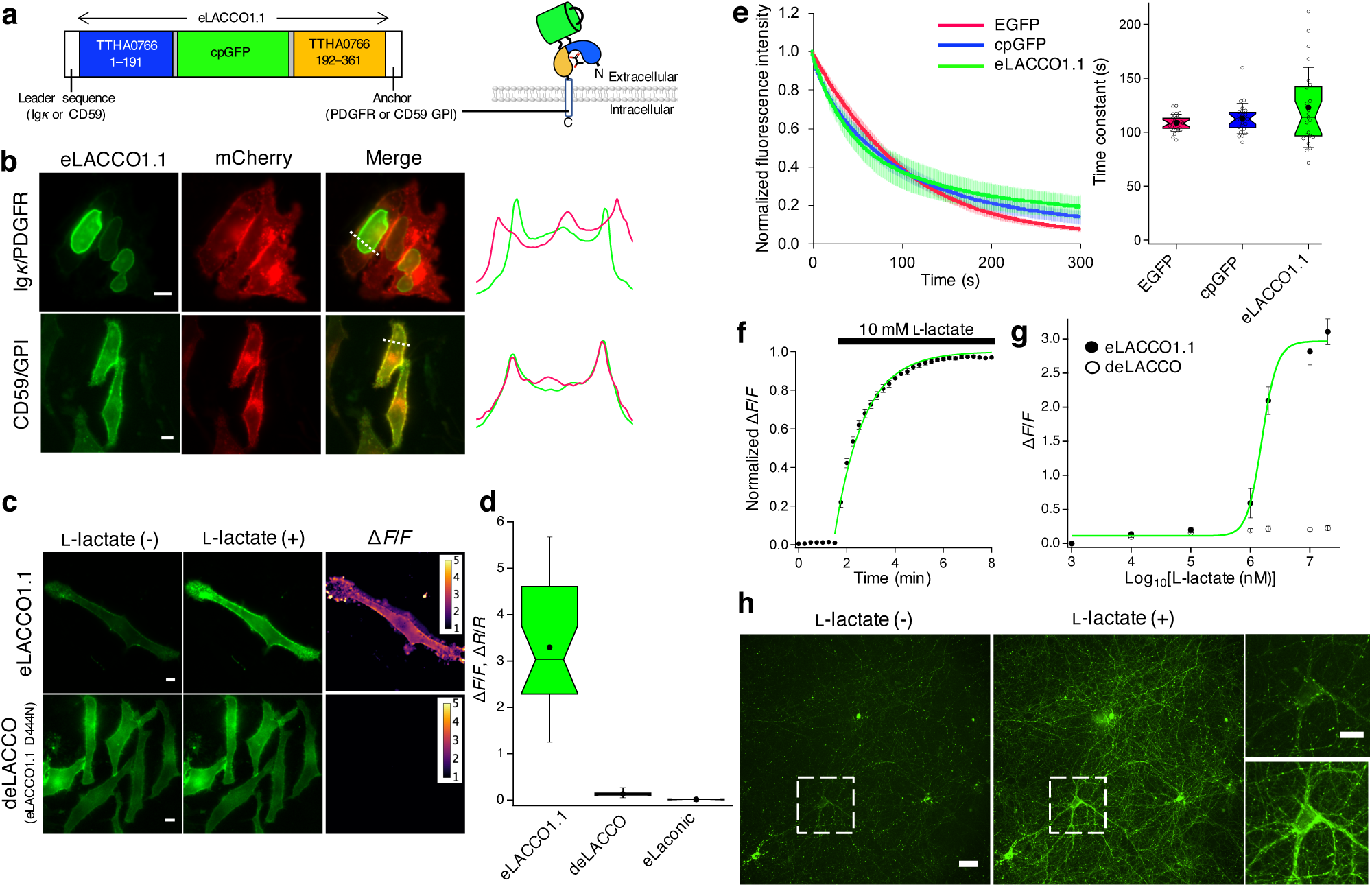
Characterization of eLACCO1.1 in live mammalian cells. (**a**) Schematic representation of the domain structure of eLACCO1.1 with N-terminal leader sequence and C-terminal anchor domain. (**b**) Localization of eLACCO1.1 with different leader sequences and anchors in HeLa cells. mCherry with Ig*κ* leader sequence and PDGFR transmembrane domain was used as cell surface marker. Line-scans (right) correspond to dashed white lines on the merged images. (**c**) Representative images of eLACCO1.1 and control biosensor deLACCO expressed on cell surface of HeLa cells before and after 10 mM L-lactate stimulation. (**d**) Box plots of L-lactate dependent Δ*F*/*F* for eLACCO1.1 and deLACCO, and Δ*R*/*R* for eLaconic. *n* = 26, 28 and 13 cells for eLACCO1.1, deLACCO and eLaconic, respectively. (**e**) Photobleaching curves (left) and time constants (right) for EGFP, cpGFP, and eLACCO1.1 expressed on the cell surface. *n* = 25, 24 and 23 cells for EGFP, cpGFP and eLACCO1.1, respectively. In the photobleaching curves, solid lines represent mean value and shaded area represent s.d. One-way ANOVA, *P* > 0.05. (**f**) Time-course of the fluorescence response of eLACCO1.1 on cell surface under L-lactate stimulation. The plot was fitted with a single exponential function. *n* = 13 cells (mean ± s.e.m.). (**g**) *In situ* titration of L-lactate on HeLa cells. Data were fitted with Hill equation. *n* = 11 cells (mean ± s.e.m.). (**h**) Representative images of eLACCO1.1 expressed in primary cortical neurons before and after 10 mM L-lactate stimulation. In the box plots of (d) and (e), the narrow part of notch with the horizontal line is the median; the top and bottom of the notch denote the 95% confidence interval of the median; the top and bottom horizontal lines are the 25^th^ and 75^th^ percentiles for the data; and the whiskers extend one standard deviation range from the mean represented as black filled circle. Scale bars in (b), (c) and (h) represent 10 μm.

To test photostability, we continuously illuminated eLACCO1.1-expressing cells using one-photon wide-field microscopy (**Fig. 2e**). eLACCO1.1 showed photostability that is comparable to EGFP and cpGFP. To examine the on-rate kinetics of eLACCO1.1, we bathed eLACCO1.1-expressing HeLa cells in a solution containing 10 mM L-lactate. L-Lactate application induced the robust increase in the fluorescence with the on rate (*τ*_on_) of 1.2 min (**Fig. 2f**). Stopped-flow analysis of purified eLACCO1.1 protein revealed that the off-rate kinetics are faster than the 1.1 ms dead time of the instrument used (**Supplementary Fig. 11**). *In situ* L-lactate titration on HeLa cells indicated that the fluorescence of eLACCO1.1 increased in a dose-dependent manner, with an apparent *K*_d_ of 1.6 mM (**Fig. 2g**). To characterize the performance of eLACCO1.1 in neurons, we expressed eLACCO1.1 in rat primary cortical neurons. eLACCO1.1 was expressed at relatively high levels, as indicated by bright membrane-localized fluorescence (**Fig. 2h**). Upon bath application of 10 mM L-lactate, eLACCO1.1 displayed the increase in fluorescence intensity with Δ*F*/*F* of 1.4 ± 0.6 (mean ± s.d. *n* > 10 neurons from 3 cultures). Taken together, these results indicated that eLACCO1.1 could potentially be useful for imaging of extracellular L-lactate concentration dynamics in living tissues.

To directly compare the performance of eLACCO1.1 with an existing L-lactate biosensor, we attempted to target the previously reported Förster resonance energy transfer (FRET)-based biosensor Laconic to the surface of HeLa cells^6^. We found that Laconic, fused with the leader and GPI anchor sequences from CD59 (designated as eLaconic), could indeed be targeted to the cell surface (**Supplementary Fig. 12a-c**). However, when using treatments similar to those used for imaging of eLACCO1.1, we were unable to observe fluorescent responses from eLaconic (**Fig. 2d**, **Supplementary Fig. 12d,e**).

### Two-photon imaging of L-lactate on astrocytes in acute brain slices

The ANLS hypothesis states that glial cells such as astrocytes can release L-lactate into the extracellular space and that this L-lactate is taken up by neurons to serve as an energy source^2^. To confirm that eLACCO1.1 remains functional on the surface of astrocytes of mammalian brain tissue, we used two-photon microscopy to examine cortical acute brain slices prepared from mice injected with an adeno-associated virus (AAV) coding eLACCO1.1 under the control of astrocyte-specific promoter (**Fig. 3a**). Bath application of L-lactate elicited a significant increase in eLACCO1.1 fluorescence at all doses tested and in a concentration-dependent manner: 1 mM (Δ*F*/*F* = 18.8 ± 4.2%, *P* = 0.01, paired *t*-test), 2.5 mM (Δ*F*/*F* = 48.0 ± 14.7%, *P* = 0.02, paired *t*-test), 10 mM (Δ*F*/*F* = 64.9 ± 16.8%, *P* = 0.01, paired *t*-test) (**Fig. 3b-d**). In contrast, the control biosensor deLACCO did not produce a significant change in fluorescence relative to baseline (Δ*F*/*F* = 0.85 ± 1.6, *P* = 0.25, paired *t*-test). Collectively, these *ex vivo* data indicate that eLACCO1.1 enables detection of extracellular L-lactate in acute brain slice and could potentially be applicable to imaging the release of L-lactate from astrocytes in this *ex vivo* brain preparation.

**Fig. 3.**
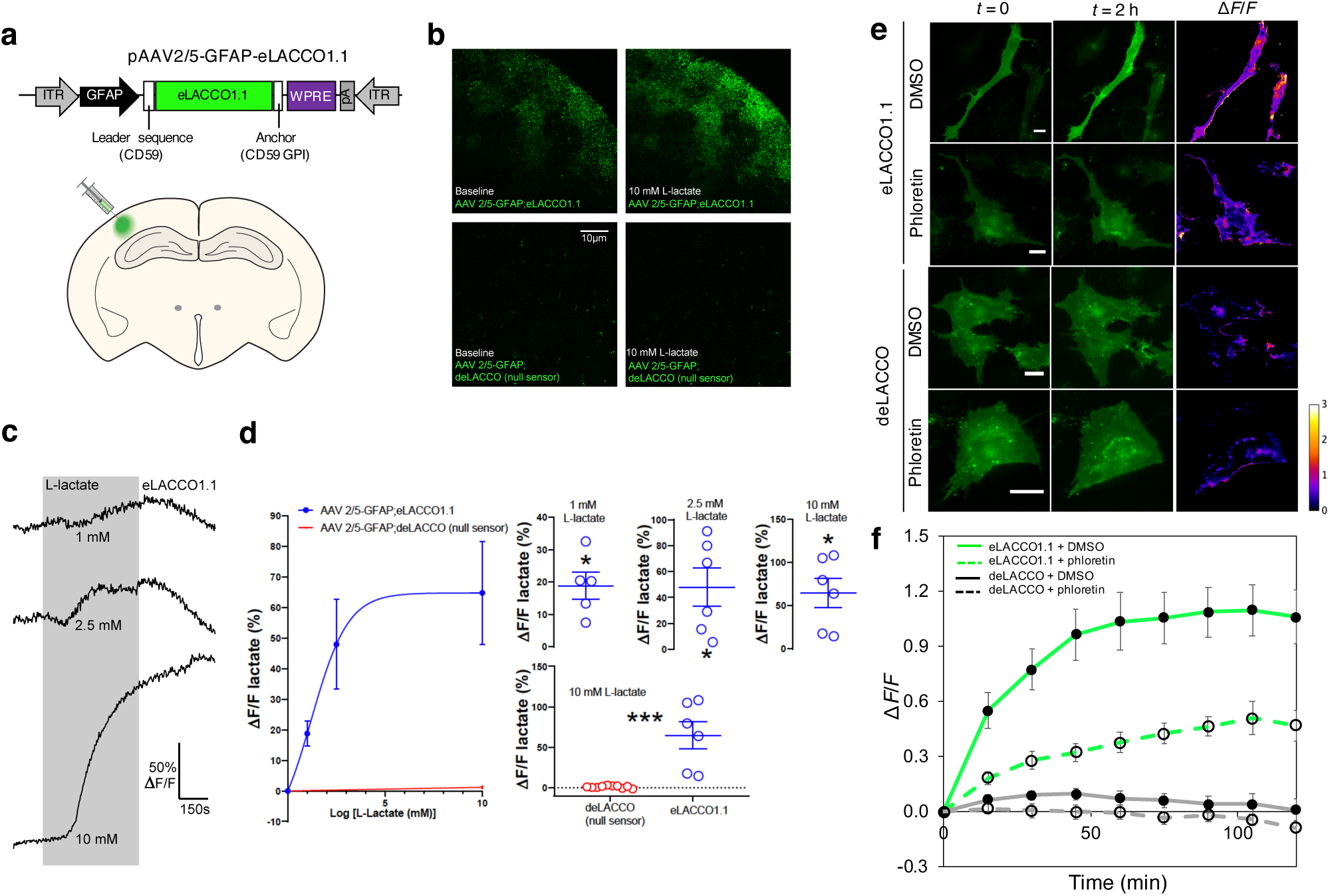
Characterization of eLACCO1.1 in acute brain slice, and imaging of endogenous L-lactate release in glioblastoma cells. (**a**) Schematic illustration of AAV injection into the somatosensory cortex for the brain slice experiments. ITR, inverted terminal repeat. GFAP, human glial fibrillary acidic protein promoter. WPRE, woodchuck hepatitis virus posttranslational regulatory element. pA, human growth hormone polyA signal. (**b**) Representative two-photon images of eLACCO1.1 and control biosensor deLACCO expressed on astrocytes in brain slice before and after 10 mM L-lactate stimulation. (**c**) Representative fluorescence traces of eLACCO1.1-expressing astrocytes in response to bath application of L-lactate. (**d**) Dose-response curves of eLACCO1.1 and deLACCO for L-lactate. eLACCO1.1; 1 mM L-lactate (n = 5 slices from 4 mice), 2.5 mM L-lactate (n = 6 slices from 4 mice), or 10 mM L-lactate (n = 6 slices from 4 mice). deLACCO; 10 mM L-lactate (n = 10 slices from 2 mice). eLACCO1.1 produced a significant increase in fluorescence upon 10 mM L-lactate relative to deLACCO control sensor (*P* < 0.001, unpaired *t*-test). Data represent mean ± s.d. (**e**) Representative fluorescence images of T98G cells expressing eLACCO1.1 or deLACCO. Scale bars, 20 μm. (**f**) Δ*F*/*F* versus time for T98G cells expressing eLACCO1.1 or deLACCO. *n* = 7 and 8 cells for eLACCO1.1 and deLACCO, respectively (mean ± s.e.m.).

### Imaging of endogenous L-lactate release from cultured glioblastoma cells

To demonstrate that eLACCO1.1 can enable imaging of the release of endogenous L-lactate from cells, we targeted eLACCO1.1 to the surface of T98G glioma cells. Under treatment with a high concentration of glucose, eLACCO1.1 localized to the surface of T98G cells underwent an increase in fluorescence consistent with increased secretion of L-lactate (**Fig. 3e,f**). In the presence of phloretin, a partial inhibitor of the monocarboxylate transporter (MCT), the glucose-dependent fluorescence increase was diminished. The control biosensor deLACCO in both the presence and absence of phloretin showed no substantial change in fluorescence intensity, indicating that the observed fluorescence changes were due to the L-lactate-dependent response of eLACCO1.1 (as opposed to a pH change or other confounding factor). Overall, these results demonstrate that eLACCO1.1 enables imaging of extracellular L-lactate release from glial cells with cellular resolution.

### Summary

This study reports the first example of a genetically encoded extracellular L-lactate biosensor, designated eLACCO1.1. We have demonstrated that eLACCO1.1 enables imaging of changes in extracellular L-lactate concentration in the context of live human cells, rat cortical neurons, and mouse brain tissue. To date, the level of extracellular L-lactate in brain tissue has typically been assessed using inserted enzyme-mediated electrodes^18^. Relative to electrodes, inherent advantages of the genetically encodable eLACCO1.1 are that it can be non-invasively introduced in the form of its corresponding gene, and it enables high resolution imaging of L-lactate concentration dynamics. In addition, the targeted expression of eLACCO1.1 in specific cell types (e.g., astrocytes or neurons), will enable researchers to accurately determine which cell types are importing, and which are exporting, L-lactate. Accordingly, we anticipate that eLACCO1.1 will prove useful for future investigations of the roles of L-lactate shuttles, including the controversial ANLS hypothesis.

## Methods

### General methods and materials

Synthetic DNA encoding the lactate binding bacterial periplasmic protein TTHA0766 was purchased from Integrated DNA Technologies. Phusion high-fidelity DNA polymerase (Thermo Fisher Scientific) was used for routine polymerase chain reaction (PCR) amplifications, and Taq DNA polymerase (New England Biolabs) was used for error-prone PCR. The QuikChange mutagenesis kit (Agilent Technologies) was used for site-directed mutagenesis. Restriction endonucleases, rapid DNA ligation kits and GeneJET miniprep kits were purchased from Thermo Fisher Scientific. PCR products and products of restriction digests were purified using agarose gel electrophoresis and the GeneJET gel extraction kit (Thermo Fisher Scientific). DNA sequences were analyzed by DNA sequence service of the University of Alberta Molecular Biology Service Unit and Fasmac Co., Ltd. Fluorescence excitation and emission spectra were recorded on a Safire2 and Spark plate reader (Tecan).

### Engineering of eLACCO1.1

The gene encoding cpGFP with N- and C-terminal linkers (LV and NP, respectively) was amplified using iGluSnFR gene as template, followed by insertion into each site of TTHA0766 lactate binding protein (LBD) in a pBAD vector (Life Technologies) by Gibson assembly (New England Biolabs). Variants were expressed in *E*. *coli* strain DH10B (Thermo Fisher Scientific) in LB media supplemented with 100 μg mL^-1^ ampicillin and 0.02% L-arabinose. Proteins were extracted using B-PER bacterial protein extraction reagent (Thermo Fisher Scientific) and tested for fluorescence brightness and lactate-dependent response. The most promising variant, designated as eLACCO0.1, was subjected to an iterative process of library generation and screening in *E*. *coli*. Libraries were generated by site-directed mutagenesis using QuikChange (Agilent Technologies) or error-prone PCR of the whole gene. For each round, 100−200 fluorescent colonies were picked, cultured and tested on 384-well plates under a plate reader. There were 9 rounds of screening before eLACCO1 was identified. Finally, Tyr80Phe mutation was added to eLACCO1 to tune the lactate affinity using Q5 high-fidelity DNA polymerase (New England Biolabs). The resulting mutant was designated as eLACCO1.1.

### Protein purification and *in vitro* characterization

The gene encoding eLACCO1.1, with a poly-histidine tag on the N terminus, was expressed from the pBAD vector. Bacteria were lysed with a cell disruptor (Branson) and then centrifuged at 15,000*g* for 30 min, and proteins were purified by Ni-NTA affinity chromatography (Agarose Bead Technologies). Absorption spectra of the samples were collected with a Lambda950 Spectrophotometer (PerkinElmer). We carried out pH titrations by diluting protein into buffers (pH from 2 to 11) containing 30 mM trisodium citrate, 30 mM sodium borate, 30 mM MOPS, 100 mM KCl, 10 mM CaEGTA, and either none or 10 mM L-lactate. Fluorescence intensities as a function of pH were then fitted by a sigmoidal binding function to determine the apparent p*K*_a_. For lactate titration, we prepared buffers by mixing a L-lactate (-) buffer (30 mM MOPS, 100 mM KCl, 1 mM CaCl_2_, pH 7.2) and a L-lactate (+) buffer (30 mM MOPS, 100 mM KCl, 1 mM CaCl_2_, 100 mM L-lactate, pH 7.2) to provide L-lactate concentrations ranging from 0 mM to 100 mM at 25°C. Fluorescence intensities were plotted against L-lactate concentrations and fitted by a sigmoidal binding function to determine the Hill coefficient and apparent p*K*_a_.

Rapid kinetic measurements of eLACCO1.1 interaction with L-lactate were made using an Applied Photophysics SX20 Stopped-flow Reaction Analyzer using fluorescence detection, exciting at 488 nm with 2 nm bandwidth and collecting emitted light at 520 nm through a 10-nm path at room temperature. The dead time of the instrument is 1.1 ms. For *k*_off_, 2 μM of purified protein sample saturated with 200 mM L-lactate and 1 mM CaCl_2_ was dissociated 1:1 with 100 mM EGTA at room temperature. Graphpad Prism was used to fit a single exponential dissociation for *k*_off_. For eLACCO1.1, *k*_off_ was faster than the dead time of the instrument, so a baseline fluorescence in the saturated state was obtained as a negative control. All measurements were done in triplicates, and error +/-represents the s.e.m.

To collect the two-photon absorption spectra, the tunable femtosecond laser InSight DeepSee (Spectra-Physics, Santa Clara, CA) was used to excite the fluorescence of the sample contained within a PC1 Spectrofluorometer (ISS, Champaign, IL). The detailed description of the optical system and experimental protocol is presented before^19^. Briefly, the laser was automatically stepped to each wavelength over the spectral range with a custom LabVIEW program (National Instruments, Austin, TX), with 42 sec at each wavelength to stabilize. We measured two samples per laser scan by using both the sample and reference holders and switching between them with the auto-switching mechanism on the PC1. The laser was focused on the sample through a 45-mm NIR achromatic lens, antireflection coating 750-1550 nm (Edmund Optics, Barrington, NJ). Fluorescence was collected from the first 0.7 mm of the sample at 90° with the standard PC1 collection optics through both 633/SP and 745/SP filters (Semrock, Rochester, NY) to remove all laser scattered light. To correct for wavelength-to-wavelength variations of laser parameters, we used LDS798 (Exciton, Lockbourne, OH) in 1:2 CHCl_3_:CDCl_3_ as a reference standard between 912-1240 nm (ref. 20), and Coumarin 540A (Exciton, Lockbourne, OH) in 1:10 DMSO:deuterated DMSO between 700-912 nm (ref. 21). Adding the deuterated solvents (Millipore Sigma, Darmstadt, Germany) was necessary to decrease NIR solvent absorption. All the dye solutions were magnetically stirred throughout the measurements. Quadratic power dependence of fluorescence intensity in the proteins and standards was checked at several wavelengths across the spectrum.

For the lactate bound state of eLACCO1.1, we measured the two-photon cross section (σ_2_) of the anionic form of the chromophore versus Rhodamine 6G in MeOH at 976 nm and 960 nm (σ_2_(976 nm) = 12.7 GM, σ_2_(960 nm) = 10.9 GM)^22^. These σ_2_ numbers closely agree with other literature data: Ref. 4 at 976 nm (considering the correction discussed in Ref. 21), and Ref. 23 at 960 nm. For the lactate free state, we measured the σ_2_ of the neutral form of the chromophore versus Fluorescein (Millipore Sigma, Darmstadt, Germany) in 10 mM NaOH (pH 12) at 820 and 840 nm (σ_2_(820 nm) = 24.2 GM, σ_2_(840 nm) = 12.9 GM^22^. These σ_2_ values for Fluorescein also match other literature data^24, 25^. Power dependence of fluorescence intensity was recorded with the PC1 monochromator at 550 nm (lactate bound) or 512 nm (lactate free) with the emission slits at a spectral width of 16 nm (FWHM) and fitted to a parabola with the curvature coefficient proportional to σ_2_. These coefficients were normalized for the concentration and the differential fluorescence quantum yield at the registration wavelength. The differential quantum yields of the standard and the sample were obtained with an integrating sphere spectrometer (Quantaurus-QY; Hamamatsu Photonics, Hamamatsu City, Japan) by selecting an integral range ∼ 8 nm to the left and right of the registration wavelength. Concentrations were determined by Beer’s Law. Extinction coefficients were determined by alkaline denaturation as detailed in Ref. 26. The two-photon absorption spectra were normalized to the σ_2_ values at the two wavelengths and averaged. To normalize to the total two-photon brightness (*F*_2_), the spectra were then multiplied by the quantum yield and the relative fraction of the respective form of the chromophore for which the σ_2_ was measured. The data is presented this way because eLACCO1.1 contains a mixture of the neutral and anionic forms of the GFP chromophore. This is described in further detail in Ref. 16 and 26.

### X-ray crystallography

For crystallization, eLACCO1 cloned in pBAD-HisB with N-terminus 6×His tagged was expressed in *E. coli* DH10B strain. Briefly, a single colony from freshly transformed *E. coli* was inoculated into 500 mL of modified Terrific Broth (2% Luria-Bertani Broth supplemented with additional 1.4% tryptone, 0.7% yeast extract, 54 mM K_2_PO_4_, 16 mM KH_2_PO_4_, 0.8% glycerol, 0.2 mg mL^-1^ ampicillin sodium salt) and shaking incubated at 37°C and 220 rpm for 8 hours to reach exponential growth phase, after which L-arabinose was added to 200 ppm to induce expression for another 48 hours shaking incubated at 30°C and 250 rpm. Bacteria were then harvested and lysed using a sonicator (QSonica) for 4 cycles of 150 sec sonication with 2 sec off between each 1 sec of sonication at 50W power. After centrifugation at 13,000*g*, the supernatant was affinity purified with Ni-NTA agarose bead (G-Biosciences) and eluted into PBS containing 250 mM imidazole and further concentrated and desalted with an Amicon Ultra-15 Centrifugal Filter Device (Merck). The Ni-NTA affinity chromatography purified eLACCO1 protein was further applied on Superdex200pg (GE healthcare) size exclusion chromatography and buffer exchanged to TBS buffer supplemented with 1 mM CaCl_2_. The fractions containing the eLACCO1 protein were pooled and concentrated to around 20 mg mL^-1^, incubated with 2 mM L-lactate for the crystallization trail. Initial crystallization of the eLACCO1 protein was set up using 384-well plate via sitting drop vapor diffusion against commercially available kits at room temperature. The eLACCO1 crystal used for the data collection was grown in 0.2 M ammonium citrate dibasic 20% w/v PEG 3350 by mixing 0.6 μl of reservoir solution with 0.6 μl of protein sample. The eLACCO1 crystals grew for 3-5 days and were cryoprotected with reservoir supplemented with 25% glycerol and frozen in liquid nitrogen. X-ray diffraction data were collected at 100 K at the Advanced Photon Source beamline line 23ID. The X-ray diffraction data were processed and scaled with XDS^27^. Data collection details and statistics were summarized in Table S1. The crystal structure of eLACCO1 was solved by molecular replacement approach implemented in Phaser program suite program^28^, using the GFP structure (PDB 3SG6) and the lactate binding domain protein (PDB 2ZZV) as search models^8, 29^. One molecules of the eLACCO1 protein was present in the asymmetric unit. Further tracing of the missing residues and the structure were iteratively rebuilt in COOT and refined with phenix.refine program^30, 31^. The final model included the most of the residues except the N terminal affinity tag and the glycine linker to connect the original N and C termini of GFP. The final eLACCO1 structure bound with one molecule of L-lactate and one Ca^2+^ ion was determined to 2.25 Å and refined to a final *R_work_/R_free_* of 0.1484/0.1871 with high quality of stereochemistry. By generating symmetry mates, the eLACCO1 packed as a dimer in the crystal packing and shared a similar dimerization interface with TTHA0766 lactate binding protein (PDB 2ZZV).

### Construction of mammalian expression vectors

For cell surface expression, the genes encoding eLACCO1.1, deLACCO, cpGFP, EGFP and Laconic (Addgene plasmid no. 44238) were amplified by PCR followed by digestion with BglII and EcoRI, and then ligated into pAEMXT vector (Covalys) that contains N-terminal leader sequence and C-terminal GPI anchor from CD59. To construct PDGFR-anchored eLACCO1.1 with various leader sequences, the gene encoding eLACCO1.1 including the CD59 leader and anchor sequence in the pAEMXT vector was first amplified by PCR followed by digestion with XhoI and HindIII, and then ligated into pcDNA3.1 vector (Thermo Fisher Scientific). Next, the gene encoding PDGFR transmembrane domain was amplified by PCR using pDisplay vector (Thermo Fisher Scientific) as a template, and then substituted with CD59 GPI domain of the pcDNA3.1 product above by using EcoRI and HindIII. Complementary oligonucleotides (Thermo Fisher Scientific) encoding each leader sequence were digested by XhoI and BglII, and then ligated into a similarly-digested pcDNA3.1 including PDGFR-anchored eLACCO1.1. The gene encoding mCherry was amplified by PCR followed by digestion with BglII and SalI, and then ligated into pDisplay vector that contains N-terminal Ig*κ* leader sequence and C-terminal PDGFR transmembrane domain. To construct eLACCO1.1 plasmid for neural expression, the gene encoding eLACCO1.1 including the CD59 leader and anchor sequence in the pAEMXT vector was first amplified by PCR followed by digestion with NheI and XhoI, and then ligated into a human synapsin promoter vector (gift from Jonathan Marvin).

### Imaging of eLACCO1.1 in HeLa, HEK293 and T98G cell lines

HeLa and HEK293FT cells were maintained in Dulbecco’s modified Eagle medium (DMEM; Nakalai Tesque) supplemented with 10% fetal bovine serum (FBS; Sigma-Aldrich) and 1% penicillin-streptomycin (Nakalai Tesque) at 37°C and 5% CO_2_. T98G cells were maintained in minimum essential medium (MEM; Nakalai Tesque) supplemented with 10% FBS, 1% penicillin-streptomycin, 1% non-essential amino acid (Nakalai Tesque) and 1 mM sodium pyruvate (Nakalai Tesque) at 37°C and 5% CO_2_. Cells were seeded in 35-mm glass-bottom cell-culture dishes (Iwaki) and transiently transfected with the constructed plasmid using polyethyleneimine (Polysciences). Transfected cells were imaged using a IX83 wide-field fluorescence microscopy (Olympus) equipped with a pE-300 LED light source (CoolLED), a 40x objective lens (numerical aperture (NA) = 1.3; oil), an ImagEM X2 EM-CCD camera (Hamamatsu), Cellsens software (Olympus) and a STR stage incubator (Tokai Hit). The filter sets used in live cell imaging had the following specification. eLACCO1.1, deLACCO, cpGFP and EGFP: excitation 470/20 nm, dichroic mirror 490-nm dclp, and emission 518/45 nm; mCherry: excitation 545/20 nm, dichroic mirror 565-nm dclp, and emission 598/55 nm; Laconic (CFP): excitation 438/24 nm, dichroic mirror 458-nm dclp, and emission 483/32 nm; Laconic (FRET): 438/24 nm, dichroic mirror 458-nm dclp, and emission 542/27 nm. Fluorescence images were analyzed with ImageJ software (National Institutes of Health).

For imaging of lactate-dependent fluorescence, HeLa cells transfected with eLACCO1.1, deLACCO, or eLaconic were washed twice with Hank’s balanced salt solution (HBSS; Nakalai Tesque), and then 2 mL of HBSS supplemented with 10 mM HEPES (Nakalai Tesque) and 10 mM 2-deoxyglucose (Wako) was added to start the imaging at 37°C followed by L-lactate stimulation. For photostability test, HeLa cells transfected with pAEMXT-eLACCO1.1, EGFP or cpGFP were illuminated by excitation light at 100% intensity of LED (∼10 mW/cm^2^ on the objective lens) and their fluorescence images were recorded at 37°C for 5 min with the exposure time of 50 ms and no interval time. Fluorescence intensities on cell membrane were collected after background subtraction. For imaging of L-lactate release, T98G cells transfected with eLACCO1.1 or deLACCO were washed twice with HBSS, and then 1.5 mL of HBSS supplemented with 10 mM HEPES was added to start the imaging in the presence of DMSO (Wako) or 100 μM phloretin (Tokyo Chemical Industry). The fluorescence images were recorded for 2 hours with the interval time of 15 min at 37°C.

### Imaging of eLACCO1.1 in neurons

Experiments were performed with rat cortical primary cultures from P0 pups, plated in glass-bottom 24-well plates. Cultures were nucleofected at time of plating, and imaged 14 days later. Three wells were plated and imaged per nucleofected construct. Culture media was replaced with 1 mL of imaging buffer (145 mM NaCl, 2.5 mM KCl, 10 mM glucose, 10 mM HEPES, 2 mM CaCl_2_, 1 mM MgCl_2_, pH 7.4) for imaging^9^. Widefield images were taken at the center of each well using a Nikon Eclipse microscope (20× 0.4 NA, 488nm excitation, 500-550nm emission) at room temperature. These were the “APO” images. After a 20-minute incubation in the presence of ∼10 mM L-lactate, the same fields of view were recorded again. These were the “SAT” images. APO and SAT images were aligned by cross-correlation. We trained a pixel classifier (Ilastik^32^) using manual annotations to label each pixel in the images as background, neurite, soma, or intracellular inclusion. A scalar constant background was subtracted from all images to account for camera offset. We report Δ*F*/*F* for pixels classified as neurites, calculated as (sum of neurite pixel intensities SAT)/(sum of neurite pixel intensities APO) - 1.

### Surgery and *in vivo* microinjections of adeno-associated virus (AAV)

Mice (C57bl/6, P45), were anesthetized via isoflurane (5% for induction, 2-3% for maintenance, v/v). Depth of anesthesia was determined by observing breathing rates and toe-pinch ensured proper loss of reflexes. Following deep anesthesia, mice were head fixed on a stereotaxic apparatus (David Kopf Instruments) with a bite bar and ear bars, with ventilated anesthesia administration. The mice were injected with 0.05 μl of buprenorphine subcutaneously (Buprenex, 0.1 mg mL^-1^), and artificial tears were applied to the eyes before beginning surgery. The hair on the scalp was removed prior to surgery, and the incision was washed with 10% povidone iodine and 70% ethanol, 3 times each, alternating. An incision was made on the scalp to expose bregma and the craniotomy site with coordinates are as follows. Somatosensory cortex: −1.58 mm posterior and +3.0 and −3.0 mm lateral (for bilateral injection) from bregma, and 0.7-0.5 mm ventrally from the pial surface. A 2-3 mm craniotomy was made at the injection site using a small burr (Fine Science Tools), powered by a drill (K.1070, Foredom). Saline (0.9%) was applied to keep the skull cool, to maintain skin hydration, and to remove bone debris. AAVs were injected via a beveled borosilicate pipette (World Precision Instruments) by a Nanoject 2 (Drummond Scientific). Five, 69 nL injections were given at each site, totaling 345 nL of virus was infused into each region of somatosensory cortex, and each virus contained the GFAP promoter driving the following constructs at the indicated titer: eLACCO1.1 (1.5×10^13^ gC mL^-1^), deLACCO (1.5×10^13^ gC mL^-1^). Following injection, the needle was left in place for 10 minutes to allow for fluid pressure normalization. Following needle withdraw, scalp was sutured with silk sutures and mice were closely monitored, kept on a heating pad and given buprenorphine twice daily for 48 hours post-op (0.05 mL, 0.1 mg mL^-1^), and fed chow with sulfonamide sulfadiazine trimethoprim (32 g kg^-1^) for 1 week post-op.

### Preparation of acute cortical brain slice

At 6 weeks after the AAV injection, mice were anaesthetized with gaseous isofluorane (5%) and then decapitated. The brain was removed, then submerged for 2 minutes in ice-cold slicing solution containing (in mM): 119.9 N-methyl-D-glucamine, 2.5 KCl, 25 NaHCO_3_, 1.0 CaCl_2_-2H_2_O, 6.9 MgCl_2_-6H_2_O, 1.4 NaH_2_PO_4_-H_2_O, and 20 D-glucose. The brain was then Krazy Glued onto a vibratome tray (Leica Instruments, VT1200S) and then re-submerged in ice-cold slicing solution. Acute coronal slices were prepared from the somatosensory cortex (400 μm thick) using a vibratome. The slices were incubated for 45 minutes at 33°C in a recovery chamber filled with artificial cerebrospinal fluid (aCSF) containing (in mM): NaCl, 126; KCl, 2.5; NaHCO_3_, 25; CaCl_2_-2H_2_O, 1.3; MgCl_2_-6H_2_O, 1.2; NaH_2_PO_4_-H_2_O, 1.25; glucose, 10. After recovery, slices were stained with astrocyte marker sulforhodamine 101 (SR101) to visualize features to maintain focal plane during imaging. Throughout, brain slices were continuously supplied with carbogen (95% oxygen, 5% carbon dioxide).

### Two-photon microscopy of eLACCO1.1 in acute cortical brain slice

Brain slices were imaged using a custom built two-photon microscope^33^ fed by a Ti:Sapphire laser source (Coherent Ultra II, ∼4 W average output at 800 nm, ∼80 MHz). Image data were acquired using MatLab (2013) running the open source scanning microscope control software ScanImage (version 3.81, HHMI/Janelia Research Campus). Imaging was performed at an excitation wavelength of 940 nm. The microscope was equipped with a primary dichroic mirror at 695 nm and green and red fluorescence was split and filtered using a secondary dichroic at 560 nm and two bandpass emission filters: 525-40 nm and 605-70 nm (Chroma Technologies). Time series images were acquired at 0.98 Hz with a pixel density of 512 by 512 and a field of view size of ∼150 μm. Imaging used a 40× water dipping objective lens (NA 1.0, WD 2.5 mm, Zeiss). Imaging was performed at room temperature. Brain slices were superfused with L-lactate (Sigma Aldrich) at concentrations of 1, 2.5, and 10 mM. Δ*F*/*F* was calculated by: Δ*F*/*F* = ((*F*_x_ – *F*_b_)/*F*_b_) × 100, where *F*_x_ is the fluorescence intensity at timepoint x and *F*_b_ is the baseline fluorescence intensity. Regions of interest were selected based on identifying fine processes via SR-101 fluorescence that did not shift focal plane during the duration of imaging.

### Animal care

For experiments performed at University of Calgary, all methods for animal care and use were approved by the University of Calgary Animal Care and Use Committee and were in accordance with the National Institutes of Health Guide for the Care and Use of Laboratory Animals. For experiments at HHMI Janelia Research Campus, all surgical and experimental procedures were in accordance with protocols approved by the HHMI Janelia Research Campus Institutional Animal Care and Use Committee and Institutional Biosafety Committee.

### Statistics and reproducibility

All data are expressed as mean ± s.d. or mean ± s.e.m., as specified in figure legends. Box plots with notches are used for Fig. 2d,e. In these plots, the narrow part of notch with the horizontal line is the median; the top and bottom of the notch denote the 95% confidence interval of the median; the top and bottom horizontal lines are the 25^th^ and 75^th^ percentiles for the data; and the whiskers extend one standard deviation range from the mean represented as black filled circle. Sample sizes (*n*) are listed with each experiment. No samples were excluded from analysis and all experiments were reproducible. In photobleaching experiments, group differences were analyzed using one-way ANOVA (Igor Pro 8). In brain slice experiments, the significant differences were analyzed using Student’s *t*-test (Graphpad Prism).

### Data availability

Structure coordinates of eLACCO1 have been deposited in the Protein Data Bank with a code of 7E9Y. Plasmids used in this study and the data that support the findings of this study are available from the corresponding author on reasonable request.

## Acknowledgments

The authors thank the University of Alberta Molecular Biology Services Unit, Y. Shen, L. Zarowny, G. Nguyen and D. Qi for technical support. We thank H. Yoshimura for the CD59 gene. Work at the University of Tokyo was supported by the Japan Society for the Promotion of Science (Grants-in-Aid for Early-Career Scientists 19K15691 and Grants-in-Aid for Scientific Research S 19H05633), Kato Memorial Bioscience Foundation, Toyota Physical and Chemical Research Institute, and The Precise Measurement Technology Promotion Foundation. Work at the University of Alberta was supported by Natural Sciences and Engineering Research Council of Canada (NSERC) and Canadian Institutes of Health Research (CIHR). Work at Montana State University was supported by National Institutes of Health (NIH) grants U01 NS094246, U24 NS109107, and F31 NS108593. Y. Wen was supported by the Alberta Parkinson Society Fellowship and National Natural Science Foundation of China (No. 31870132 and N0. 82072237).

## Author Contributions

Y.N. developed eLACCO1.1 and performed *in vitro* characterization. R.S.M. and M.D. measured one-photon absorbance spectra and two-photon excitation spectra. C.M.R. and J.H. performed acute brain slice imaging. S.Z. and Y.W. worked on the crystallography of eLACCO1. A.A. performed stopped-flow experiment. A.A. and K.P. performed the imaging of primary neurons and data analysis. Y.K. performed screening of leader sequence. M.-E.P produced AAV. M.J.L., K.P., J.S.B., G.R.G. and R.E.C. supervised research. Y.N. and R.E.C. wrote the manuscript.

## Supplementary information

**Supplementary Figure 1.**
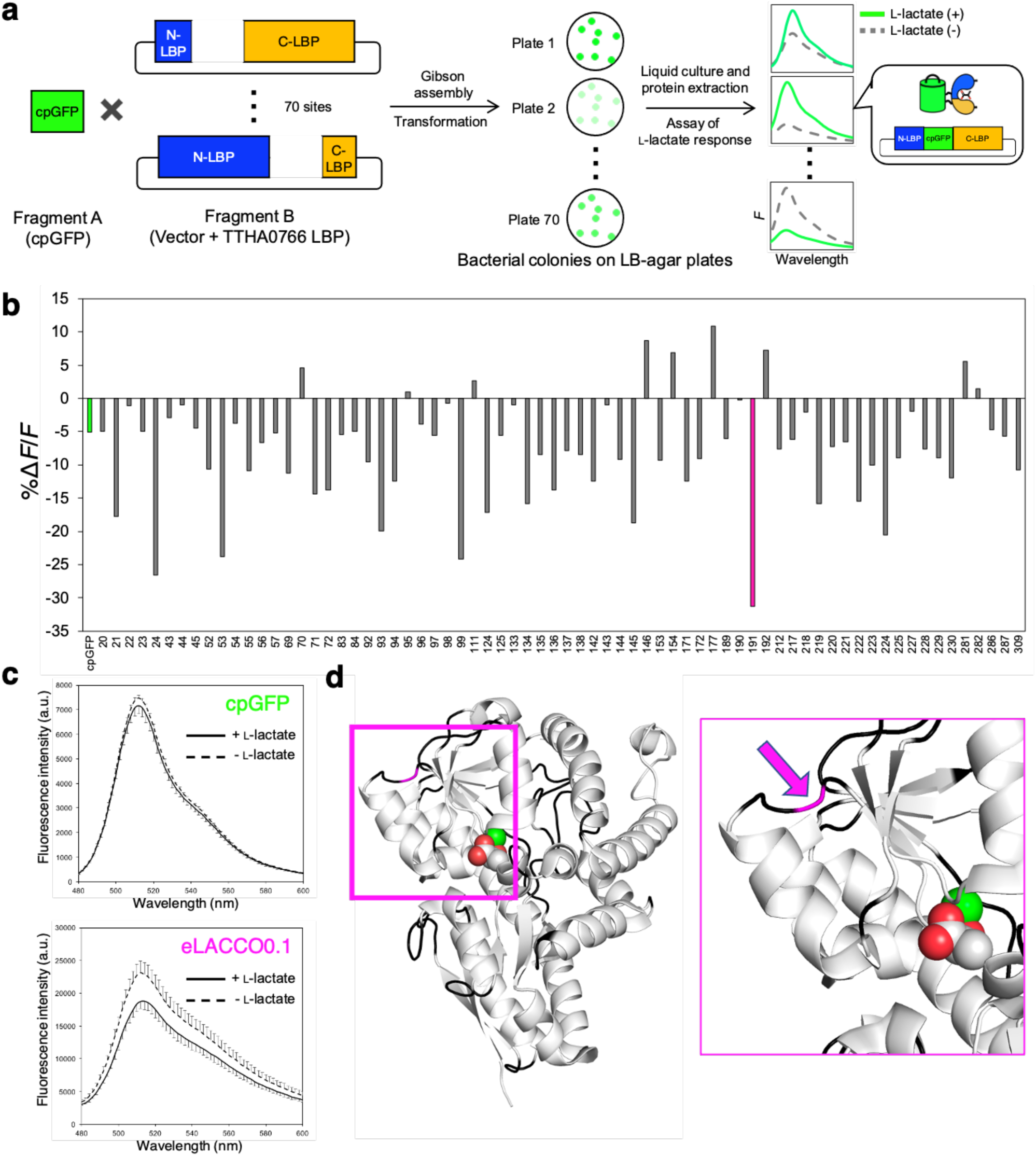
Construction of the biosensor prototype. (**a**) Schematic illustration of the biosensor prototype construction. TTHA0766 L-lactate binding protein (LBP) genes with a range of split site were ligated with cpGFP gene by Gibson assembly. Transformation of *E. coli* with assembled plasmids resulted in colonies expressing each biosensor prototype protein. Fluorescent colonies were cultured followed by extraction of the protein, and then the change in fluorescence intensity upon 10 mM L-lactate (Δ*F*/*F*) was tested. (**b**) Δ*F*/*F* profile of each candidate protein for prototype biosensor. The number of horizontal axis represents the cpGFP insertion site on TTHA0766. Δ*F*/*F* of cpGFP as a negative control is shown in green. Red bar at the insertion site 191 showed the largest absolute value of Δ*F*/*F* and the protein was designated eLACCO0.1. (**c**) Emission spectra of cpGFP and eLACCO0.1 in the presence and absence of 10 mM L-lactate. Error bars represent standard deviation of triplicates. (**d**) Crystal structure of TTHA0766 (PDB ID 2ZZV). All insertion sites tested are colored in black. The insertion site 191 is highlighted in magenta.

**Supplementary Figure 2.**
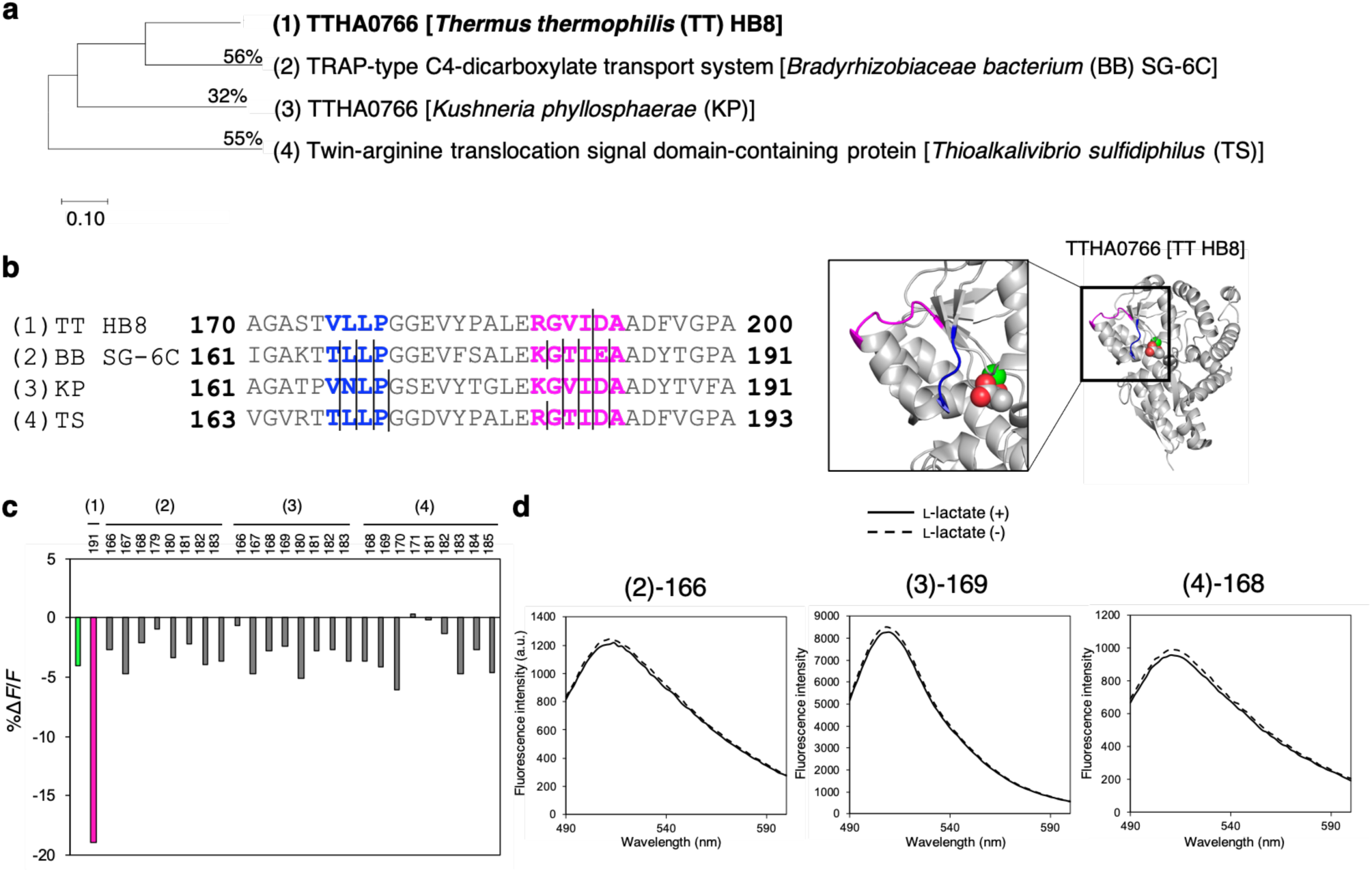
Sensor prototypes based on the various TTHA0766 homologues. (**a**) Phylogenic tree of the various TTHA0766 homologues. Sequence identities of the whole gene in amino acid with *Thermus thermophilis* TTHA0766 are shown as percentages. (**b**) Alignment of the four sequences in the vicinity of the insertion sites. Predicted loop regions from the crystal structure of *Thermus thermophilis* TTHA0766 (right) are highlighted in blue and magenta. Lines represent the sites for cpGFP insertion. (**c**) Δ*F*/*F* profile of each candidate protein for prototype biosensor. The numbers along the horizontal axis represent the cpGFP insertion site on each target protein. Δ*F*/*F* of cpGFP as a negative control and eLACCO0.1 are colored in green and magenta, respectively. (**d**) Representative emission spectra of the prototype biosensor proteins in the presence and absence of 10 mM L-lactate.

**Supplementary Figure 3.**
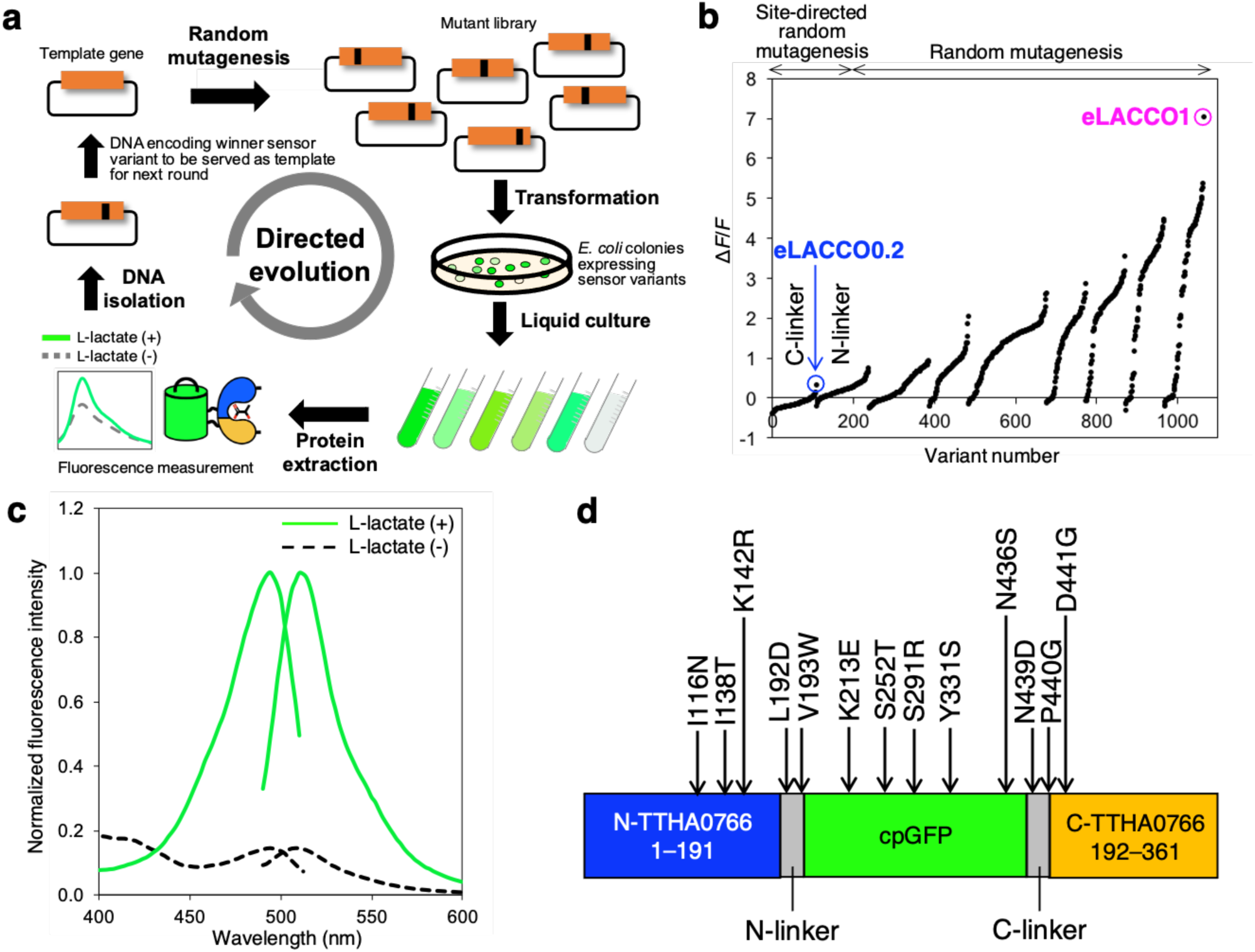
Directed evolution of eLACCO1. (**a**) Schematic of directed evolution workflow. Specific sites (i.e., the linkers) or the entire gene of template L-lactate biosensor genes were randomly mutated and the resulting mutant library was used to transform *E. coli*. Bright colonies were picked and cultured, and then proteins were extracted to examine Δ*F*/*F* upon addition of 10 mM L-lactate. The variant with the highest Δ*F*/*F* was used as the template for the next round. (**b**) Δ*F*/*F* rank plot representing all proteins tested during the directed evolution. For each round, tested variants are ranked from lowest (negative responses indicate inverse response) to highest Δ*F*/*F* value from left to right. Nine rounds of the evolution led to eLACCO1 indicated with a magenta circle. Inset highlights the plot of the first round by random mutagenesis of C-terminal linker. One variant, designated eLACCO0.2, showed the direct fluorescence response to L-lactate while the template eLACCO0.1 decreased the fluorescence intensity (inverse response) in response to L-lactate. (**c**) Excitation and emission spectra of eLACCO1 in the presence and absence of 10 mM L-lactate. (**d**) Mutations accumulated during the directed evolution to arrive at eLACCO1.

**Supplementary Figure 4.**
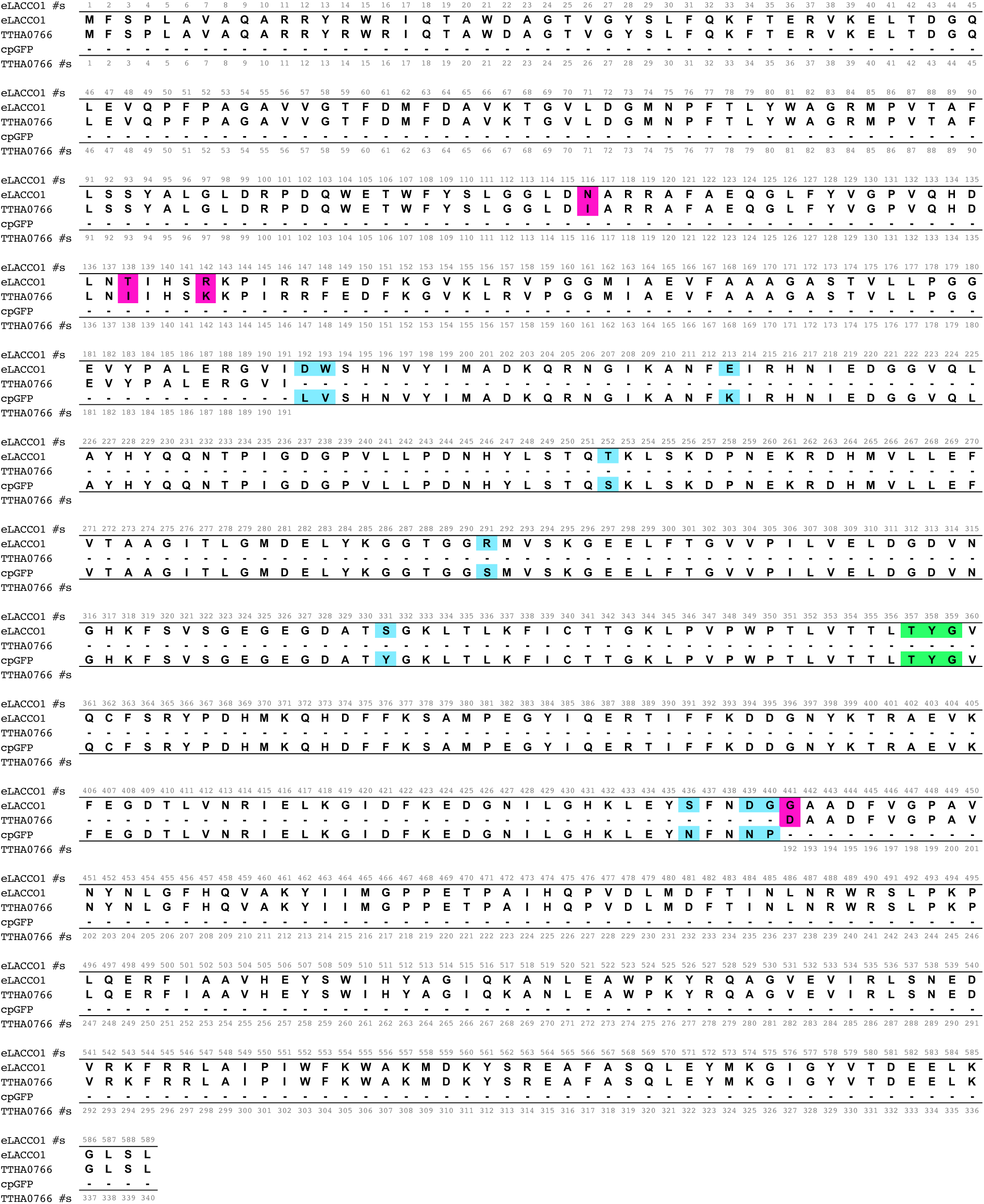
Sequence alignment of TTHA0766, cpGFP, and eLACCO1. Mutations in eLACCO1 (**refer to Supplementary Figure 3d**), relative to TTHA0766 and cpGFP, are highlighted in magenta and blue, respectively. The chromophore-forming residues are highlighted in green.

**Supplementary Figure 5.**
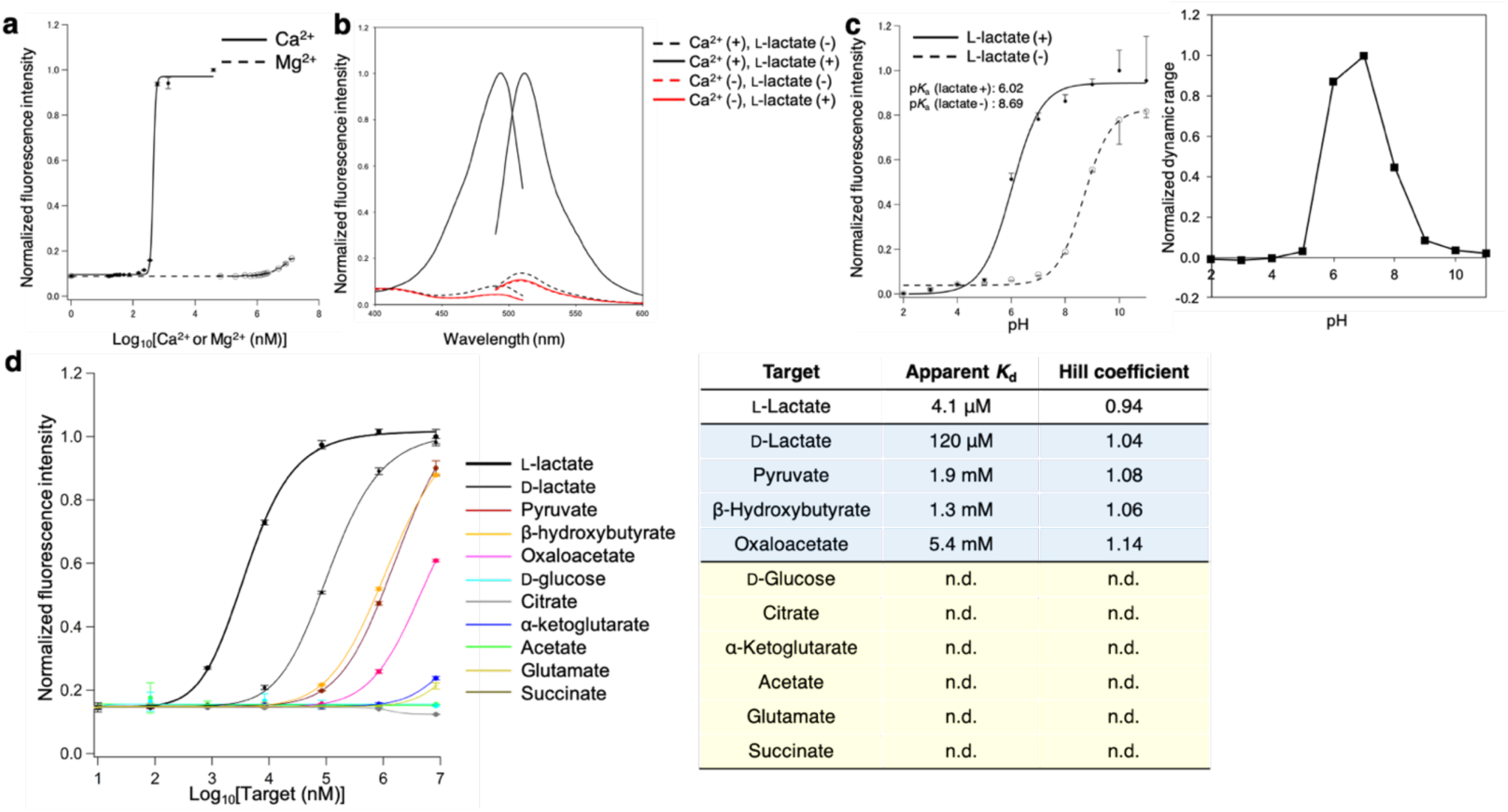
*In vitro* characterization of eLACCO1. (**a**) Fluorescence of eLACCO1 in the presence of 10 mM L-lactate as a function of Ca^2+^ and magnesium ion (Mg^2+^). *n* = 3 independent experiments (mean ± s.d.). (**b**) Excitation and emission spectra of eLACCO1 in the presence and absence of 10 mM L-lactate and 39 μM Ca^2+^. (**c**) pH titration curves of eLACCO1 in the presence and absence of 10 mM L-lactate. *n* = 3 independent experiments (mean ± s.d.). (**d**) Dose-response curves of eLACCO1 for L-lactate and a variety of metabolites. *n* = 3 independent experiments (mean ± s.d.). Right table summarizes apparent *K*_d_ and Hill coefficient for each target molecule. n.d., not determined.

**Supplementary Figure 6.**
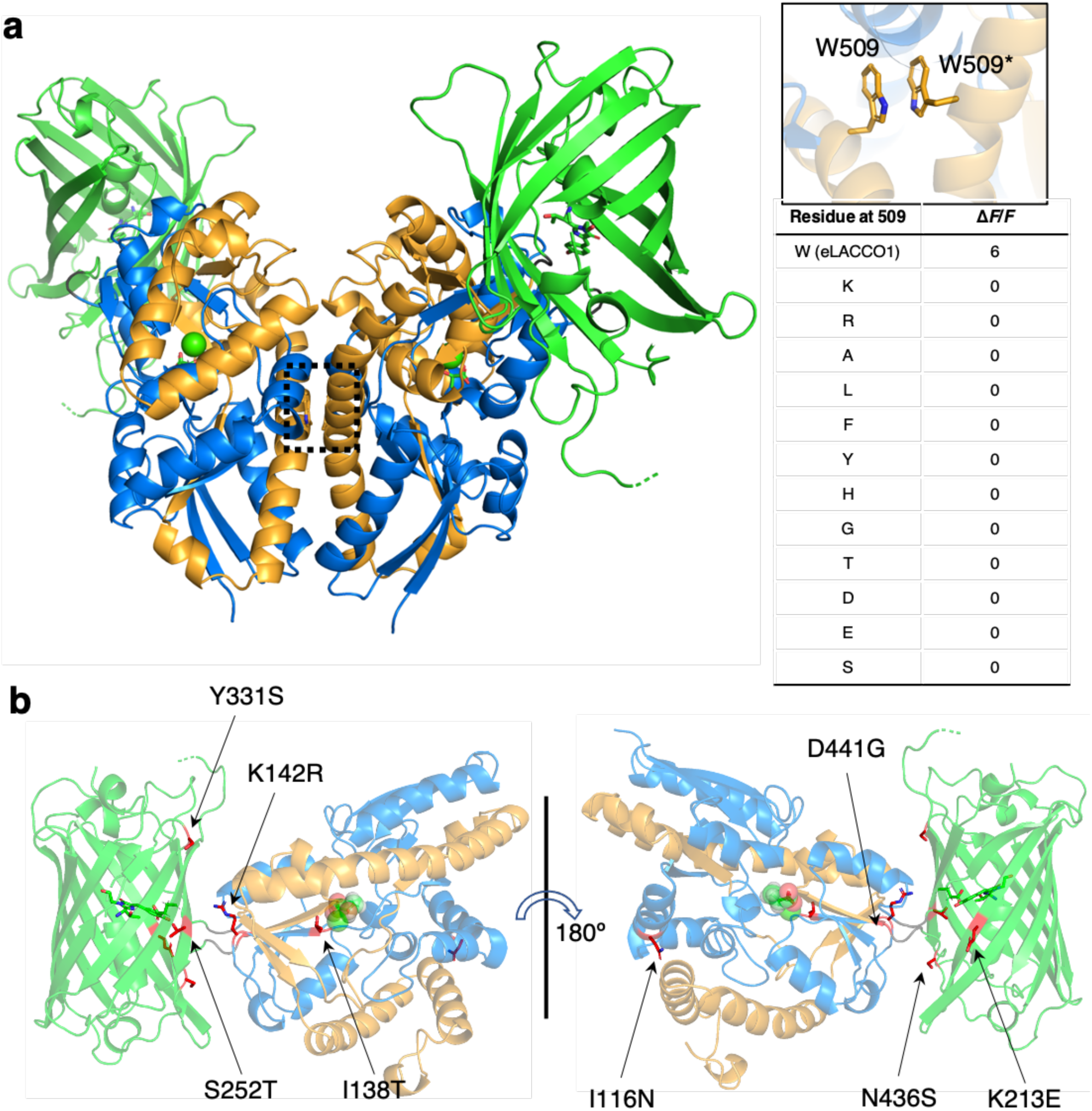
Crystal structure of eLACCO1. (**a**) Dimer structure of eLACCO1. W509 residues of each protomer form a hydrophobic interaction via π-π stacking. In an effort to disrupt the dimer interface and make the protein monomeric, we performed the site-directed mutagenesis on W509. L-Lactate is represented in stick format. Green sphere represents Ca^2+^. Table summarizes Δ*F*/*F* of W509 variants. (**b**) Overall representation of the eLACCO1 crystal structure with the position of mutations indicated. L-Lactate is shown in a sphere representation.

**Supplementary Figure 7.**
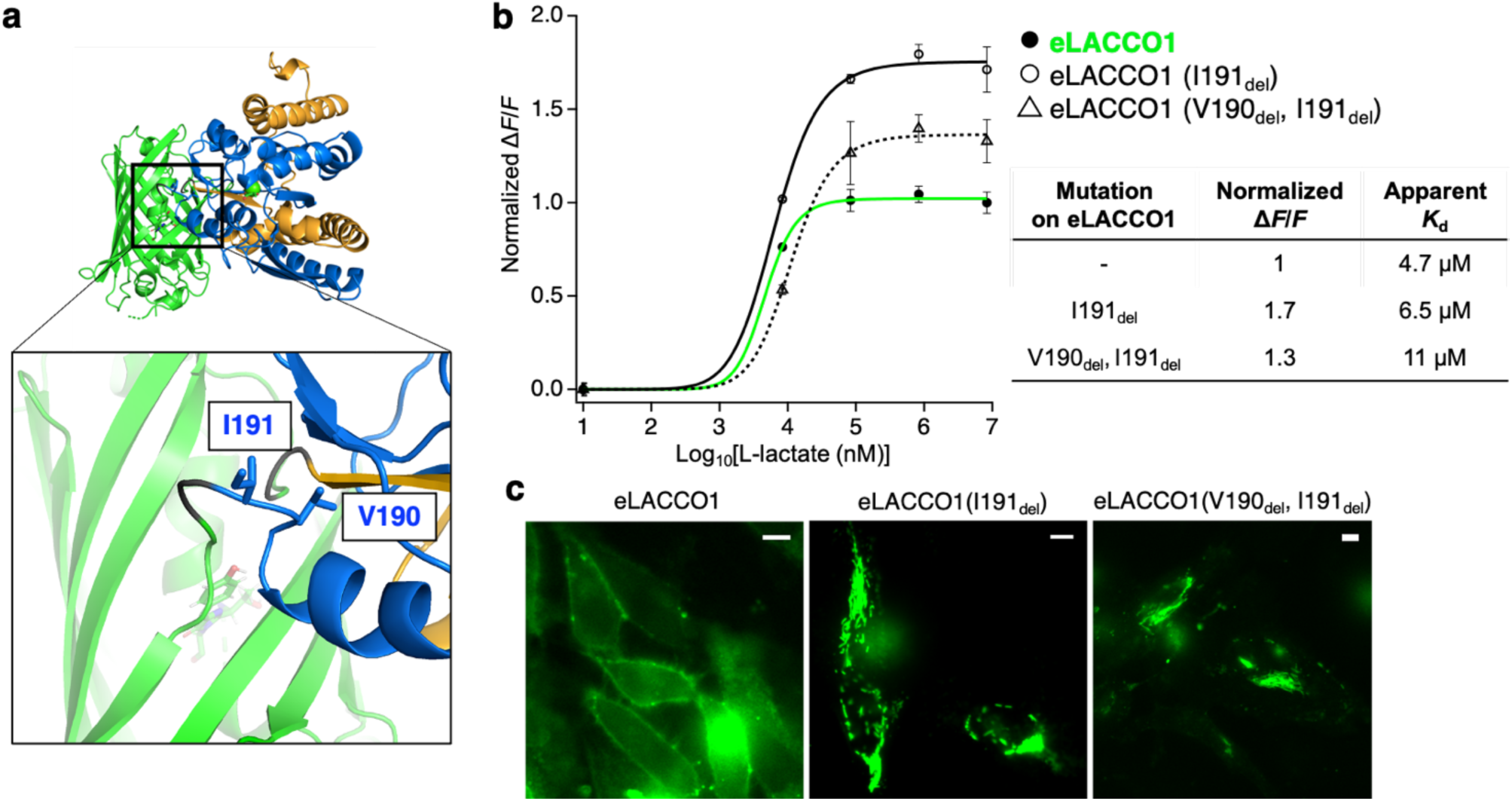
Optimization of the linker between cpGFP and TTH00766 in eLACCO1. (**a**) Zoom-in view of the region around N-terminal linker of eLACCO1. (**b**) Dose-response curves of eLACCO1 and its deletion variants for L-lactate. *n* = 3 independent experiments (mean ± s.d.). Right table summarizes the normalized Δ*F*/*F* and apparent *K*_d_ for each protein. (**c**) Localization of eLACCO1 and its deletion variants in living HeLa cells. Scale bars, 10 μm.

**Supplementary Figure 8.**
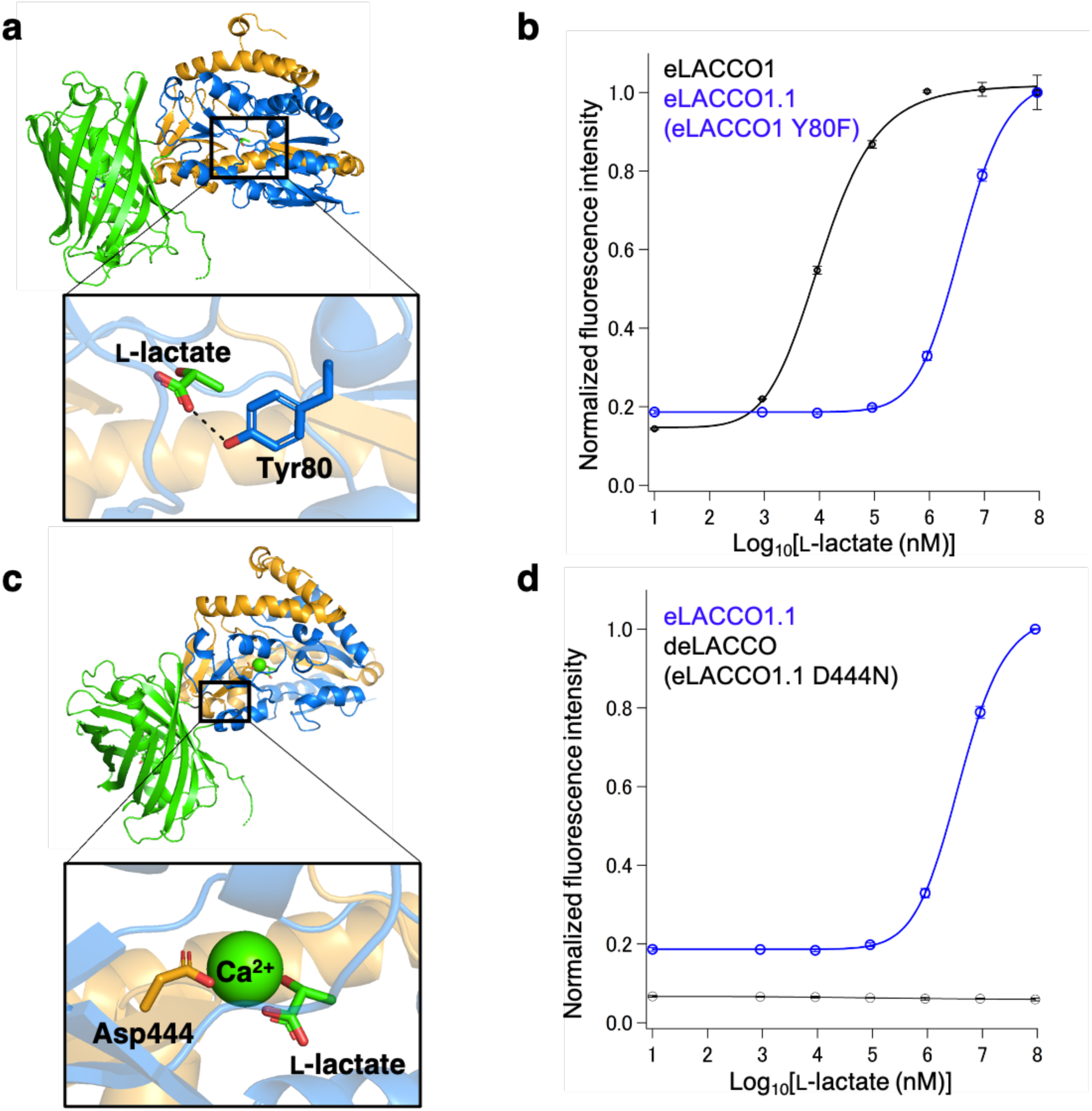
Affinity tuning of eLACCO1. (**a**) Crystal structure of eLACCO1 and zoom-in view of the L-lactate binding pocket. The phenol group of the Tyr80 side chain forms a hydrogen bond to the carboxylate group of L-lactate. (**b**) Fluorescence of eLACCO1 and eLACCO1.1 (eLACCO1 Y80F) as a function of L-lactate. (**c**) Crystal structure of eLACCO1 and zoom-in view of the Ca^2+^ binding pocket. Carboxyl group of Asp444 side chain coordinates to Ca^2+^. (**d**) Fluorescence of eLACCO1.1 and deLACCO (eLACCO1.1 D444N) as a function of L-lactate.

**Supplementary Figure 9.**
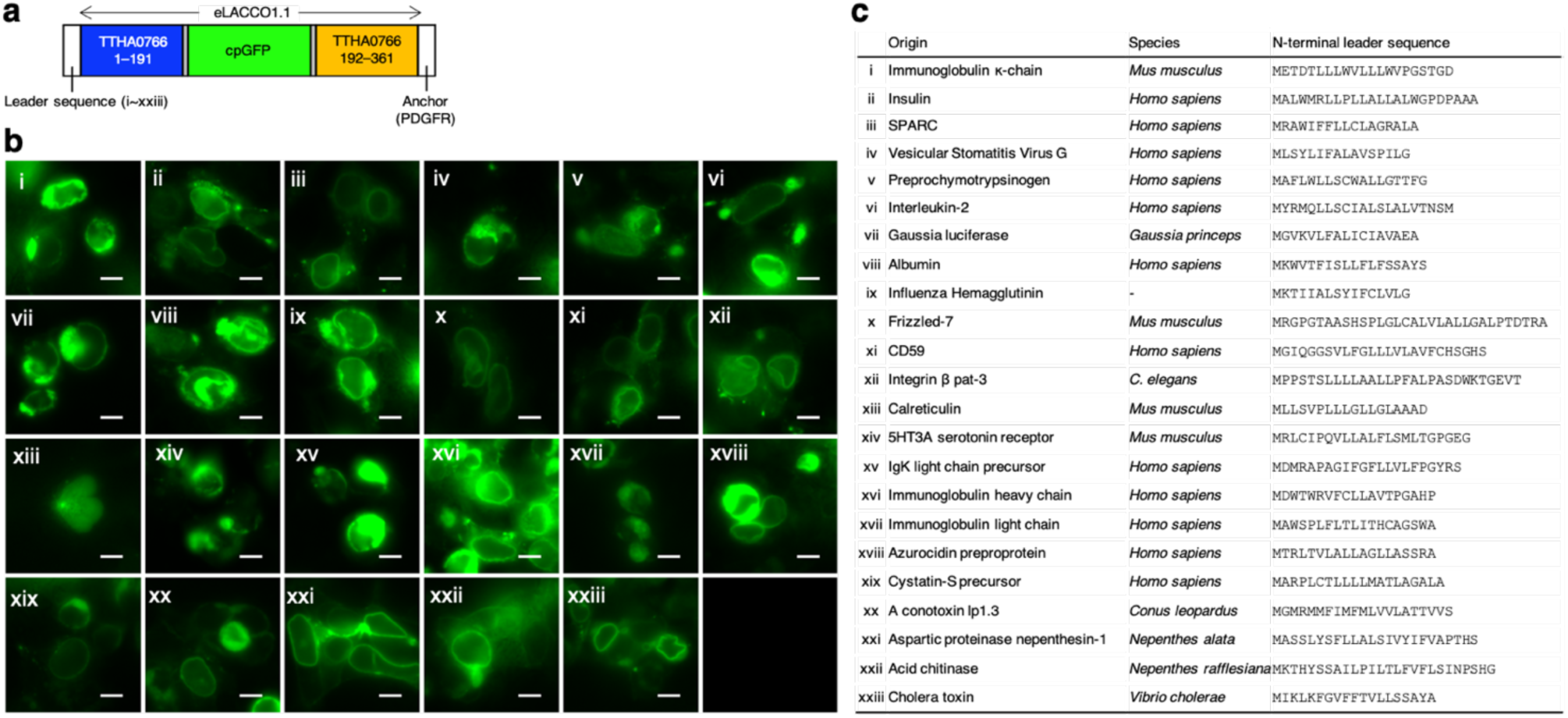
Membrane trafficking of eLACCO1.1 with various leader sequences. (**a**) Schematic of the primary sequence of eLACCO1.1 with N-terminal leader sequence and C-terminal PDGFR transmembrane anchor. (**b**) Localization of PDGFR-anchored eLACCO1.1 with different leader sequences in HEK293FT cells. Scale bars, 10 μm. (**c**) Amino acid sequences of the leader peptide tested.

**Supplementary Figure 10.**
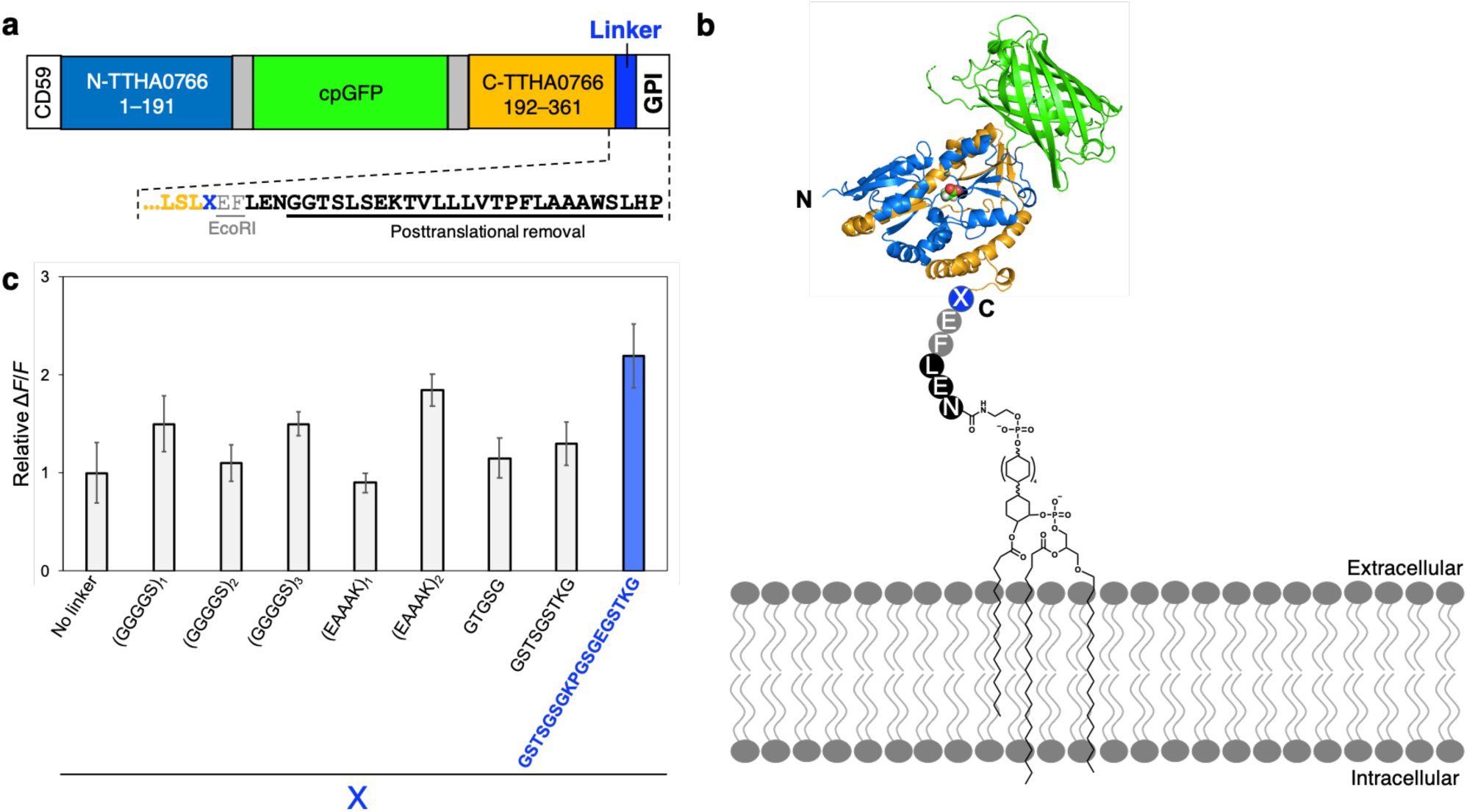
Optimization of the linker between eLACCO1.1 and GPI anchor. Schematic of the primary sequence of eLACCO1.1 with CD59 leader sequence and CD59 GPI anchor. Underlined sequence is removed after the translation, followed by GPI labeling to the C-terminal Asn(N) residue. (a) Schematic representation of GPI-anchored eLACCO1.1 on cell surface. (**c**) Relative Δ*F*/*F* of a range of eLACCO1.1 linker variants. The linker GSTSGSGKPGSGEGSTKG has been proven to reduce the aggregation and be resistant to proteolysis^1^. *n* = 6 cells (mean ± s.e.m.).

**Supplementary Figure 11.**
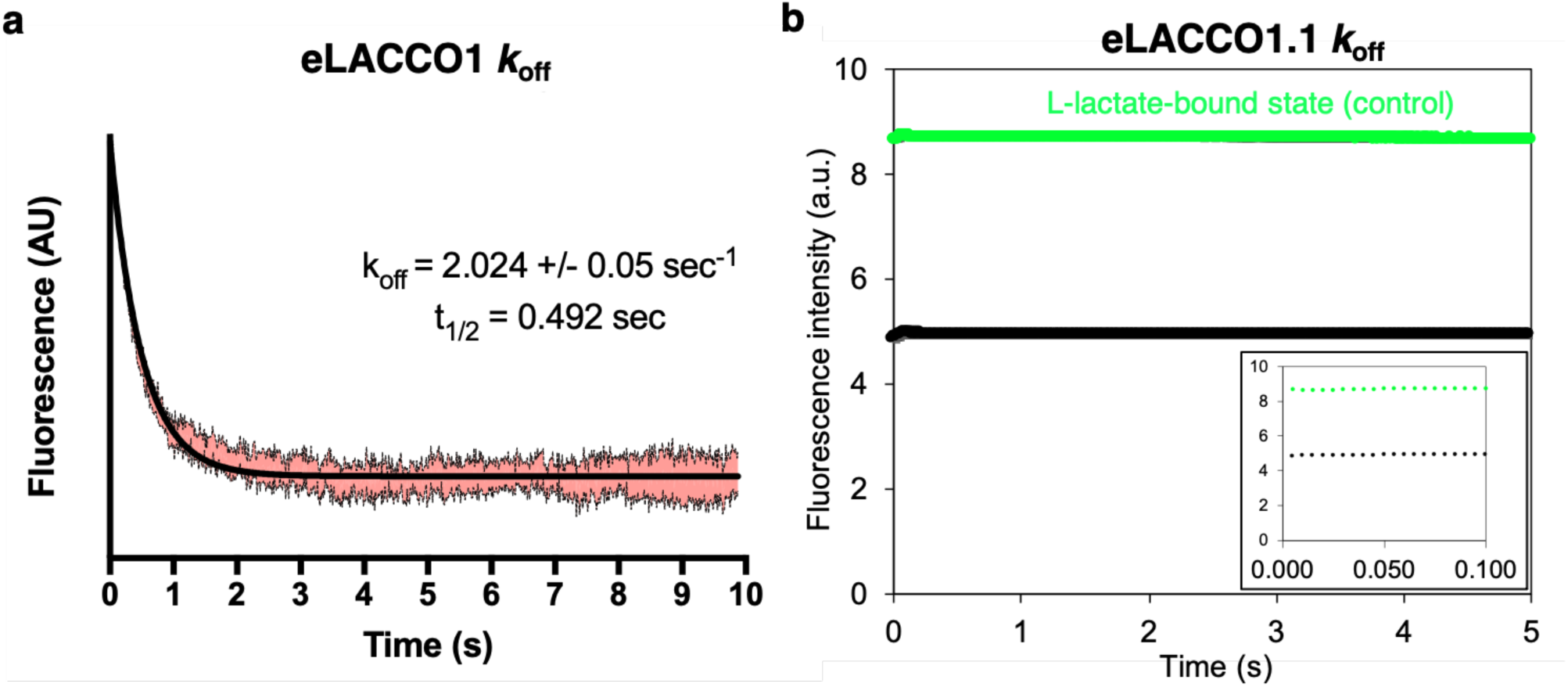
Stopped-flow analysis of eLACCO1 and eLACCO1.1. (**a**) Stopped-flow characterization of purified eLACCO1. L-Lactate-free medium was rapidly mixed at t = 0. (**b**) Stopped-flow characterization of purified eLACCO1.1. L-Lactate-free medium was rapidly mixed at t = 0 (black filled circle). Green filled circles represent the fluorescence intensity of L-lactate-bound eLACCO1.1 as a negative control. Inset is the same data at the time < 0.1 s.

**Supplementary Figure 12.**
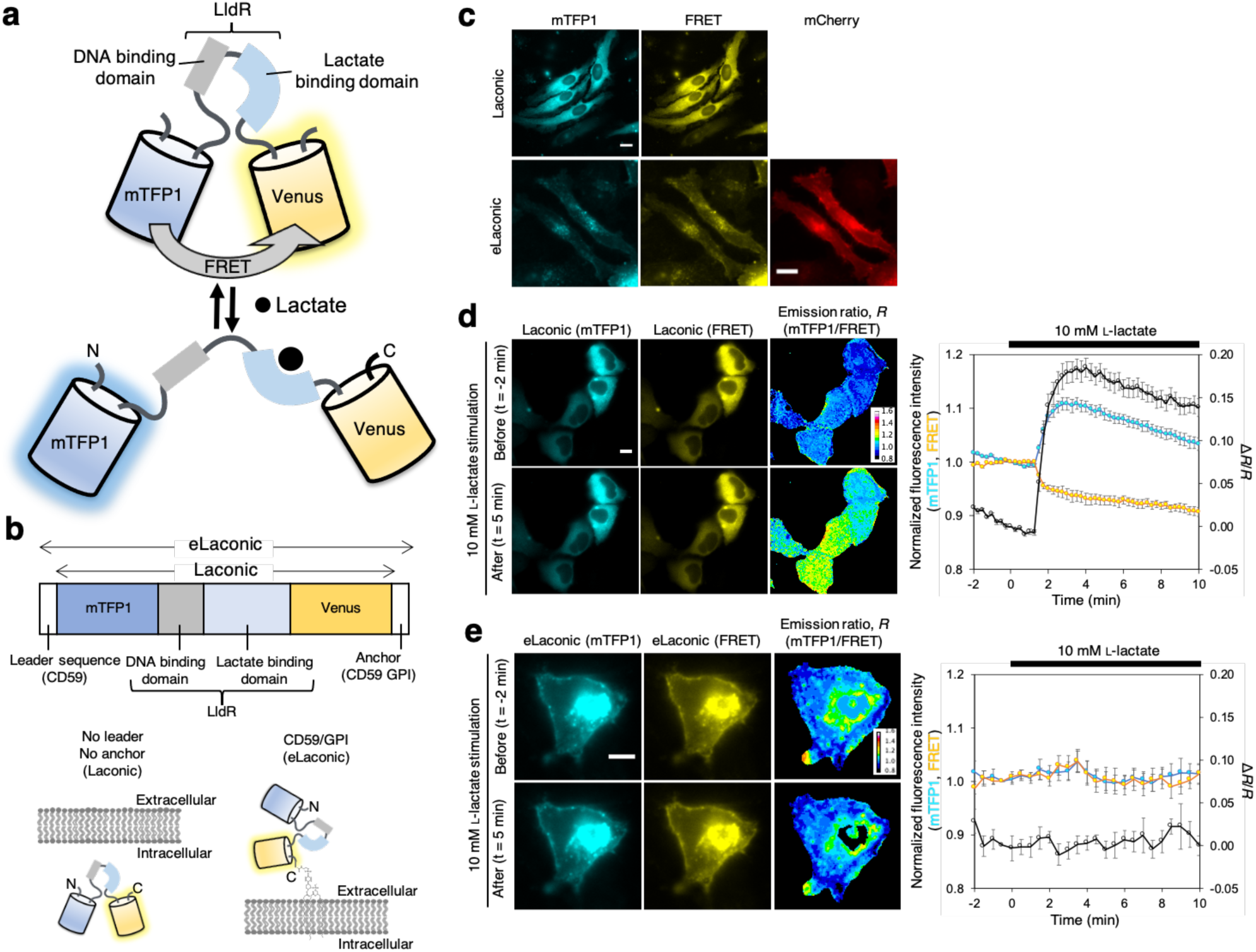
Live cell imaging with Laconic. (**a**) Schematic representation of Laconic. Binding of L-lactate to LldR L-lactate/DNA binding domain decreases FRET efficiency, thereby changing the fluorescence intensities of mTFP1 cyan fluorescent protein and Venus yellow fluorescent protein^2^. (**b**) Schematic of the primary sequence of Laconic. We named Laconic with N-terminal leader sequence and C-terminal anchor domain as eLaconic. (**c**) Representative images of HeLa cells expressing Laconic or eLaconic. mCherry with Ig*κ* leader sequence and PDGFR transmembrane domain was used as cell surface marker. Scale bars, 20 μm. (**d**) Representative images of HeLa cells expressing Laconic before and after 10 mM L-lactate stimulation. HeLa cells were pre-incubated for 1 hour with iodoacetic (500 μM) to block glycolysis at GAPDH, and then imaged in the presence of H^+^/K^+^ exchanger nigericine (10 μM) and rotenone (2 μM) to equilibrate the extracellular and intracellular L-lactate^2^. Right graph represents the time course of the fluorescence intensity (mTFP1 and FRET) and Δ*R*/*R*, where *R* is the emission ratio (mTFP1/FRET). *n* = 8 cells (mean ± s.e.m.). Scale bar, 10 μm. (**e**) Representative images of HeLa cell expressing eLaconic before and after 10 mM L-lactate stimulation. Right graph represents the time course of the fluorescence intensity (mTFP1 and FRET) and Δ*R*/*R*. *n* = 13 cells (mean ± s.e.m.). Scale bar, 10 μm.

**Supplementary Table 1.** Crystallographic and refinement statistics of eLACCO1.

| Crystal | eLACCO1 |
| --- | --- |
| **Data collection** |  |
| Space group | C222<sub>1</sub> |
| *a*, *b*, *c* (Å) | 102.2, 105.0, 138.7 |
| $\alpha$, $\beta$, $\gamma$ (°) | 90.0, 90.0, 90 |
| Resolution (Å) | 52.49-2.25 (2.33-2.25) |
| *R*<sub>merge</sub> | 0.132 (1.14) |
| *R*<sub>meas</sub> | 0.158 (1.3) |
| Multiplicity | 5.3 (5.2) |
| CC(1/2) | 0.986 (0.57) |
| CC* | 0.997 (0.75) |
| *I*/$\sigma$(*I*) | 5.75 (1.0) |
| Completeness (%) | 99.93 (99.97) |
| Wilson B-factor (Å<sup>2</sup>) | 19.33 |
| **Refinement** |  |
| Total Reflections | 187301 (18505) |
| Unique Reflections | 35639 (3530) |
| *R*<sub>work</sub>/*R*<sub>free</sub> | 0.1484/0.1871 (0.1557/0.2109) |
| **Number of atoms** |  |
| Protein | 4497 |
| Ligands | 29 |
| Water | 511 |
| Average B-factor (Å<sup>2</sup>) | 21.66 |
| Protein ADP (Å<sup>2</sup>) | 20.87 |
| Ligands (Å<sup>2</sup>) | 17.46 |
| Water | 28.87 |
| **Ramachandran plot** |  |
| Favored/Allowed (%) | 97/3 |
| **Root-Mean-Square-Deviations:** |  |
| Bond lengths (Å) | 0.012 |
| Bond Angle (°) | 1.44 |
| PDB code | 7E9Y |

Statistics for the highest resolution shell are shown in parentheses.

**Supplementary Table 2.**
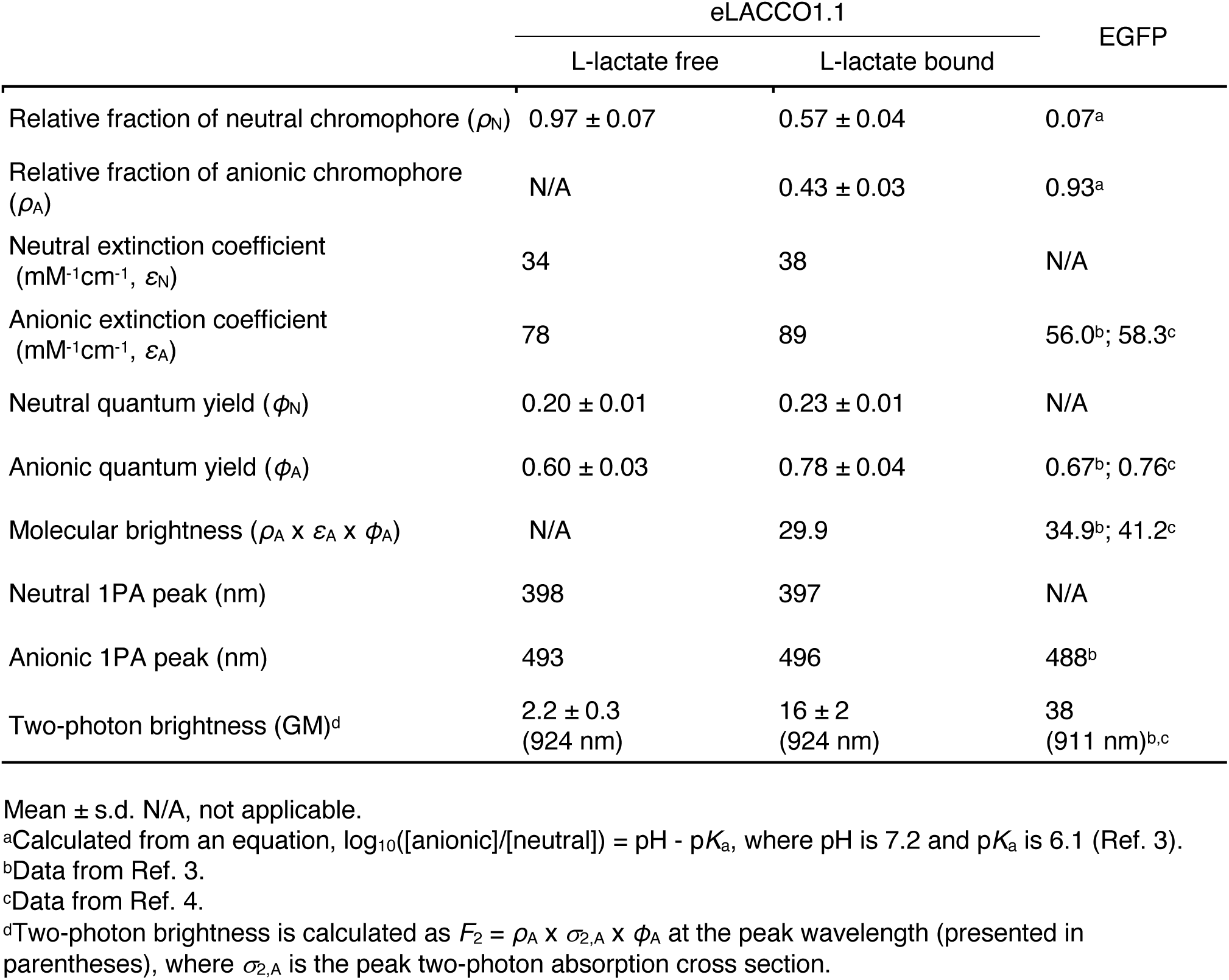
One-photon and two-photon photophysical parameters of eLACCO1.1.

## References

1. Rabinowitz, J., Enerbäck, S. Nat. Metab. 2, 566–571 (2020).

2. Magistretti, P., Allaman, I. Nat. Rev. Neurosc*i.* 19, 235–249 (2018).

3. Yellen, G. J. Cell Biol. 217, 2235–2246 (2018).

4. Greenwald, E. C., Mehta, S., Zhang, J. Chem. Rev. 118, 11707–11794 (2018).

5. Nasu, Y., Shen, Y., Kramer, L. & Campbell, R. E. Nat. Chem. Biol. https://doi.org/10.1038/s41589-020-00718-x (2021).

6. San Martin, A., et al. PLoS ONE 8, e57712 (2013).

7. Harada, K. et al. Sci. Rep. 10, 19562 (2020).

8. Akiyama, N., Takeda, K., Miki, K., J. Mol. Biol. 392, 559–565 (2009).

9. Chen, T.-W et al. Nature 499, 295–300 (2013).

10. Murphy-Royal, C. et al. Nat. Commun. 11, 2014 (2020).

11. Ferguson, B. et al. Eur. J. Appl. Physiol. 118, 691–728 (2018).

12. Chattoraj, M., King, B. A., Bublitz, G. U., & Boxer, S. G. Proc. Natl Acad. Sci. USA, 93, 8362–8367 (1996).

13. Cranfill, P. et al. Nat. Methods 13, 557–562 (2016).

14. Molina, R. S., J. Phys. Chem. Lett. 8, 2548–2554 (2017).

15. Drobizhev, M. et al. Nat. Methods 8, 393–399 (2011).

16. Molina, R. et al. Biophys. J. 116, 1873–1886 (2019).

17. Yang, L. et al. Anesth. Analg. 105, 673–679 (2007).

18. Zuend, M. et al. Nat. Metab. 2, 179–191 (2020).

19. Drobizhev, M., Molina, R. S., & Hughes, T. E. Bio-Protocol, 10, https://doi.org/10.21769/BioProtoc.3498 (2020).

20. Makarov, N.S., Campo, J., Hales, J.M. & Perry, J.W. Opt. Mater. Express, OME 1, 551–563 (2011).

21. de Reguardati, S., Pahapill, J., Mikhailov, A., Stepanenko, Y. & Rebane, A. Opt. Express 24, 9053–9066 (2016).

22. Makarov, N. S., Drobizhev, M. & Rebane, A. Opt. Express, OE 16, 4029–4047 (2008).

23. Albota, M. A., Xu, C. & Webb, W. W. Appl. Opt. 37, 7352–7356 (1998).

24. Xu, C. & Webb, W. W. J. Opt. Soc. Am. B, 13, 481–491 (1996).

25. Rodriguez, L., Ahn, H.-Y. & Belfield, K. D. Opt. Express 17, 19617–19628 (2009).

26. Barnett, L. M., Hughes, T. E. & Drobizhev, M. PLoS One 12, e0170934 (2017).

27. Klotz, J.; Porter, B. E.; Colas, C.; Schlessinger, A.; Pajor, A. M. Mol. Med. 22, 310−321 (2016).

28. A. J. Mccoy, et al. J. Appl. Crystallogr. 40, 658–674 (2007).

29. Akerboom, J. et al. J. Neurosci. 32, 13819–13840 (2012).

30. Emsley, P., Lohkamp, B., Scott, W. G. & Cowtan, K. Acta Crystallogr. D Biol. Crystallogr. 66, 486–501 (2010).

31. Adams, P. D. et al. Acta Crystallogr. D Biol. Crystallogr. 66, 213–221 (2010).

32. Berg, S. et al. Nat. Methods 16, 1226–1232 (2019).

33. Rosenegger D. G. et al. PLoS One 9, e110475 (2014).

## Supplementary References

1. Whitlow, M. et al. Protein Eng. DesSel 8, 989–995 (1993).

2. San Martin, A. et al. PLoS ONE 8, e57712 (2013).

3. Cranfill, P. et al. Nat. Methods 13, 557–562 (2016).

4. Molina, R. S. et al. J. Phys. Chem. Lett. 8, 2548–2554 (2017)

